# Head-direction cells escaping attractor dynamics in the parahippocampal region

**DOI:** 10.1101/268110

**Authors:** Olga Kornienko, Patrick Latuske, Laura Kohler, Kevin Allen

## Abstract

Navigation depends on the activity of head-direction (HD) cells. Computational models postulate that HD cells form a uniform population that reacts coherently to changes in landmarks. We tested whether this applied to HD cells of the medial entorhinal cortex and parasubiculum, areas where the HD signal contributes to the periodic firing of grid cells. Manipulations of the visual landmarks surrounding freely-moving mice altered the tuning of HD cells. Importantly, these tuning modifications were often non-coherent across cells, refuting the notion that HD cells form a uniform population constrained by attractor-like dynamics. Instead, examination of theta rhythmicity 1revealed two types of HD cells, theta rhythmic and non-rhythmic cells. Larger tuning alterations were observed predominantly in non-rhythmic HD cells. Moreover, only non-rhythmic HD cells reorganized their firing associations in response to visual landmark changes. These findings reveal a theta non-rhythmic HD signal whose malleable organization is controlled by visual landmarks.

## Introduction

Efficient navigation in mammals depends on several classes of spatially selective neurons. The animal’s sense of direction is encoded by head-direction (HD) cells that track ongoing angular movements of the head (Taube et al., 1990a; Valerio and Taube, 2012; Butler et al., 2017). Each HD cell fires maximally when the head of an animal points in a particular direction, with different cells having different preferred directions. HD cells are located in both subcortical and cortical areas including the dorsal tegmental nucleus, lateral mammillary nucleus, anterodorsal thalamic nucleus, postsubiculum, retrosplenial cortex, parasubiculum, and the medial entorhinal cortex (Taube et al., 1990a; Chen et al., 1994; Taube, 1995; Stackman and Taube, 1998; Sharp et al., 2001; Sargolini et al., 2006; Cacucci et al., 2004). The HD signal is generated in subcortical areas, possibly in the dorsal tegmental and lateral mammillary nuclei, and depends on intact vestibular inputs (Stackman and Taube, 1997; Muir et al., 2009; Yoder and Taube, 2009; Valerio and Taube, 2016). This signal reaches cortical areas via the thalamocortical excitatory projections of the anterodorsal thalamic nucleus. Accordingly, lesions to the subcortical components of the HD system strongly impairs HD representations in cortical areas (Goodridge and Taube, 1997; Bassett et al., 2007; Winter et al., 2015).

Continuous attractor network models provide a mechanistic framework explaining how the HD signal is generated (Skaggs et al., 1995; Redish et al., 1996; Zhang, 1996). In these models, neurons are allotted positions in a circle according to their preferred firing direction. The connectivity between neurons depends on their relative position. While neighboring cells with similar preferred HD excite each other, distant cells tend to inhibit each other. This connectivity leads to the emergence of an activity packet which represents the moment-to-moment HD of the animal. These models predict that preferred HD differences between HD cells never change because the connections within the network are immutable.

Most empirical data support attractor network models of HD cells. For example, rotation of a peripheral visual landmark in an environment causes an equivalent rotation of the preferred direction of all HD cells (Taube et al., 1990b). When distal and proximal cues on a circular maze are rotated in opposite directions, HD cells in the thalamus rotate coherently (Yoganarasimha et al., 2006). Additional evidence comes from recordings of HD cells after manipulating the vestibular system. After occlusion of the semicircular canals, HD cells become unstable relative to external landmarks (Muir et al., 2009). Nevertheless, the temporal firing order of HD cell pairs remains dependent on the animal’s turning direction, suggesting that HD cells drift coherently, as predicted. The tendency of HD cells to fire together has also been compared in different brain states (Peyrache et al., 2015). As predicted by the models, firing associations of HD cells observed during exploratory behavior are maintained during sleep. Further more, reliable spike transmission is observed between HD cells in the anterodorsal thalamic nucleus and the presubiculum (Peyrache et al., 2015), suggesting that cortical cells inherit their preferred HD from thalamic HD cells.

The medial entorhinal cortex and parasubiculum (MEC/PaS) are located at the top of hierarchical ascending HD pathways (Clark and Taube, 2012). There, HD cells intermingle with speed cells and grid cells (Sargolini et al., 2006; Boccara et al., 2010; Kropff et al., 2015). It has been proposed that a key function of these cells is to compute an estimate of the animal’s position in space, which is optimally represented by the activity of modularly organized grid cells (McNaughton et al., 2006; Stensola et al., 2012; Herz et al., 2017). To keep track of the animal’s position during movement, the grid cell network is thought to integrate direction and speed of movement using the activity of HD and speed cells, respectively (McNaughton et al., 2006; Sargolini et al., 2006; Kropff et al., 2015). Thus, this spatial code likely depends on the properties of its HD inputs (Winter et al., 2015).

The MEC/PaS receive important visual and visuospatial inputs from the postrhinal and retrosplenial cortices (Wyss and Van Groen, 1992; Burwell and Amaral, 1998; Czajkowski et al., 2013). Indeed, recent studies have established that a substantial fraction of MEC/PaS neurons encode information about visual patterns or contexts (Pérez-Escobar et al., 2016; Diehl et al., 2017; Ismakov et al., 2017). Whether and how visual information impacts the HD signal in the MEC/PaS is not known. One possibility is that visual landmarks only contribute to setting and stabilizing the preferred direction of HD cells. This scenario is consistent with current computational models and with recordings performed in the anterodorsal thalamic nucleus and the presubiculum (Skaggs et al., 1995; Redish et al., 1996; Zhang, 1996; Stackman et al., 2003; Yoder et al., 2011). Alternatively, some HD cells might encode information about both the HD and the visual landmarks, causing HD cells to react non-coherently to manipulations of the environment and therefore escape the attractor-like dynamics predicted by HD cell models.

We tested this alternative hypothesis by recording the activity of HD cells in the MEC/PaS of mice exploring an open-field environment. This environment was comprised of an elevated platform surrounded by four walls. Two distinct light patterns located on different walls were turned on and off. We report that changing which visual pattern was presented caused changes in the tuning curves of the HD cells. Simultaneous recordings from multiple HD cells revealed that the tuning changes in HD cells were often non-coherent, demonstrating that the activity of some HD cells is not constrained by attractor-like dynamics. Further analysis showed that HD cells could be divided into two classes: theta rhythmic and non-rhythmic HD cells. Pronounced landmark-driven non-coherent tuning changes were observed in non-rhythmic HD cells. In contrast, theta rhythmic HD cells maintained their firing associations.

## Results

To investigate the impact of visual landmarks on the HD signal, we recorded the activity of MEC/PaS neurons together with oscillatory field potentials using multichannel extracellular techniques in freely-behaving mice. Nine mice were trained to explore an elevated square platform surrounded by 4 walls (Figure 1a). Two distinct visual patterns (vp1 and vp2) made from different arrangements of LED strips were attached to two other adjacent walls (Figure 1a). The two distinct visual patterns were switched on and off in turn. Each recording session included forty 2-min trials alternating between vp1 and vp2 (Figure 1b). In addition, a standard cue card attached to one wall remained at the same location throughout the experiment (Figure 1a).

**Figure 1.**
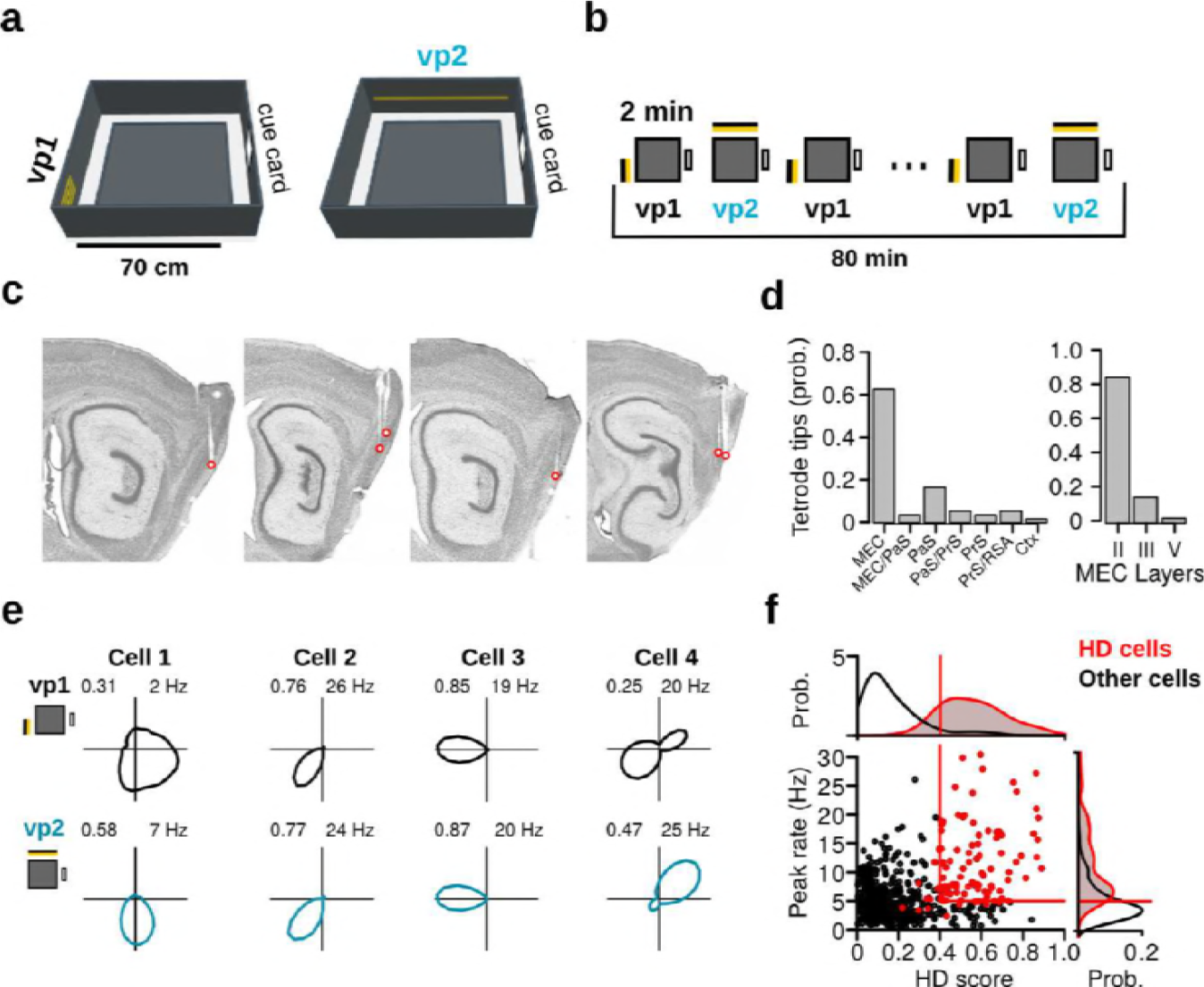
Recording protocol, histological results and examples of HD cells. **a**, The recording environment was an elevated square platform surrounded by four walls. Two distinct visual patterns (vp1 and vp2) made of LED strips were attached to two adjacent walls. A standard paper cue card was attached to a third wall. **b**, Recording sessions comprised a sequence of forty 2-min trials that alternated between vp1 and vp2 trials. **c**, Sagittal brain sections showing representative recording sites in the MEC and PaS. Red circles indicate tetrode tips. **d**, Distribution of tetrode tips across brain regions and different layers of the MEC. PaS: parasubiculum, PrS: Presubiculum, RSA: retrosplenial agranular cortex, Ctx: cortex. **e**, HD firing rate polar plots for four HD cells recorded during the two light conditions (numbers indicate HD score and peak firing rate). **f**, Scatter plot showing HD scores and peak firing rates of all neurons during vp2 trials. Each dot represents one cell. Lines indicate thresholds for HD cells identification. Red dots are HD cells.

Histological analysis revealed that most recording sites were in the MEC (65.4%, 34 out of 52; Figures 1c, 1d and Supplementary Figure 1; Supplementary Table 1). Half of the remaining recording sites were found in the PaS (17.3%, 9 out of 52). Of the recording sites in the MEC, 82.4% (28 out of 34) had entered layer II of the MEC before the end of the experiment (Figure 1d).

A total of 944 neurons were recorded over 167 recording sessions. The HD tuning curve of each neuron was calculated separately for trials with vp1 and vp2 (Figure 1e). The HD score, which was defined by the mean vector length of the tuning curve, was used as a measure of HD selectivity. Cells with a HD score exceeding 0.4, a peak firing rate larger than 5 Hz, and a directional distributive ratio larger than 0.2 during the vp1 or vp2 trials were considered putative HD cells (104 out of 944 neurons, Figure 1f). The 3 directional distributive ratio ensured that HD selectivity was not a byproduct of spatial selectivity coupled with unequal HD sampling across the recording environment (Muller et al., 1994; Cacucci et al., 2004) (see Materials and Methods and Supplementary Figure 2a-c). Out of 104 putative HD cells, 10.5% (*N* = 11) were conjunctive grid x HD cells. These were excluded from the HD cell category, leaving a total of 93 HD cells. Of the remaining HD cells, 29.0% were speed modulated, 61.3% had significant spatial sparsity scores, and 31.2% were HD selective only (Supplementary Figure 2d).

We confirmed that the neurons expressed similar levels of directional tuning during the vp1 and vp2 trials. At the population level, HD scores, peak firing rates and mean firing rates of HD cells were not significantly different between the two trial types (Supplementary Figure 2f). Preferred directions were randomly distributed during both trial types (Rayleigh test of uniformity, vp1: test statistic = 0.0931, *P* = 0.4466; vp2: test statistic = 0.0479, *P* = 0.81). Moreover, the behavior of mice was comparable between the two different trial types (Supplementary Figure 2g).

### Visual landmarks alter the tuning Curves of HD Cells

We investigated whether the two visual patterns caused modifications in the tuning curves of HD cells. Inspection of the tuning curves during vp1 and vp2 trials suggested that the preferred direction of some HD cells changed with the different visual patterns (Figure 2a). For some HD cells, these changes could be observed between consecutive trials, causing the preferred direction of the neurons to oscillate in time (Figure 2a middle). To test whether the change of preferred direction in each cell was statistically significant, we compared the observed change to a surrogate distribution of changes obtained by randomly reassigning trial labels (vp1 and vp2) (Figure 2a right, see Materials and Methods). Observed changes that were larger than 99% of the surrogate changes were considered significant. We found that 63.4% (59 out of 93) of the HD cells significantly altered their preferred HD between vp1 and vp2 trials.

A similar analysis was performed to test whether the level of directionality of HD cells changed with the visual patterns presented to the animal (Figure 2b). We observed clear examples of HD cells that changed their selectivity depending on the visual patterns (Figure 2b). These changes could often be observed between single trials (Figure 2b middle). Overall, 47.3% (44 out of 93) of the HD cells significantly altered their HD score between trial types.

**Figure 2.**
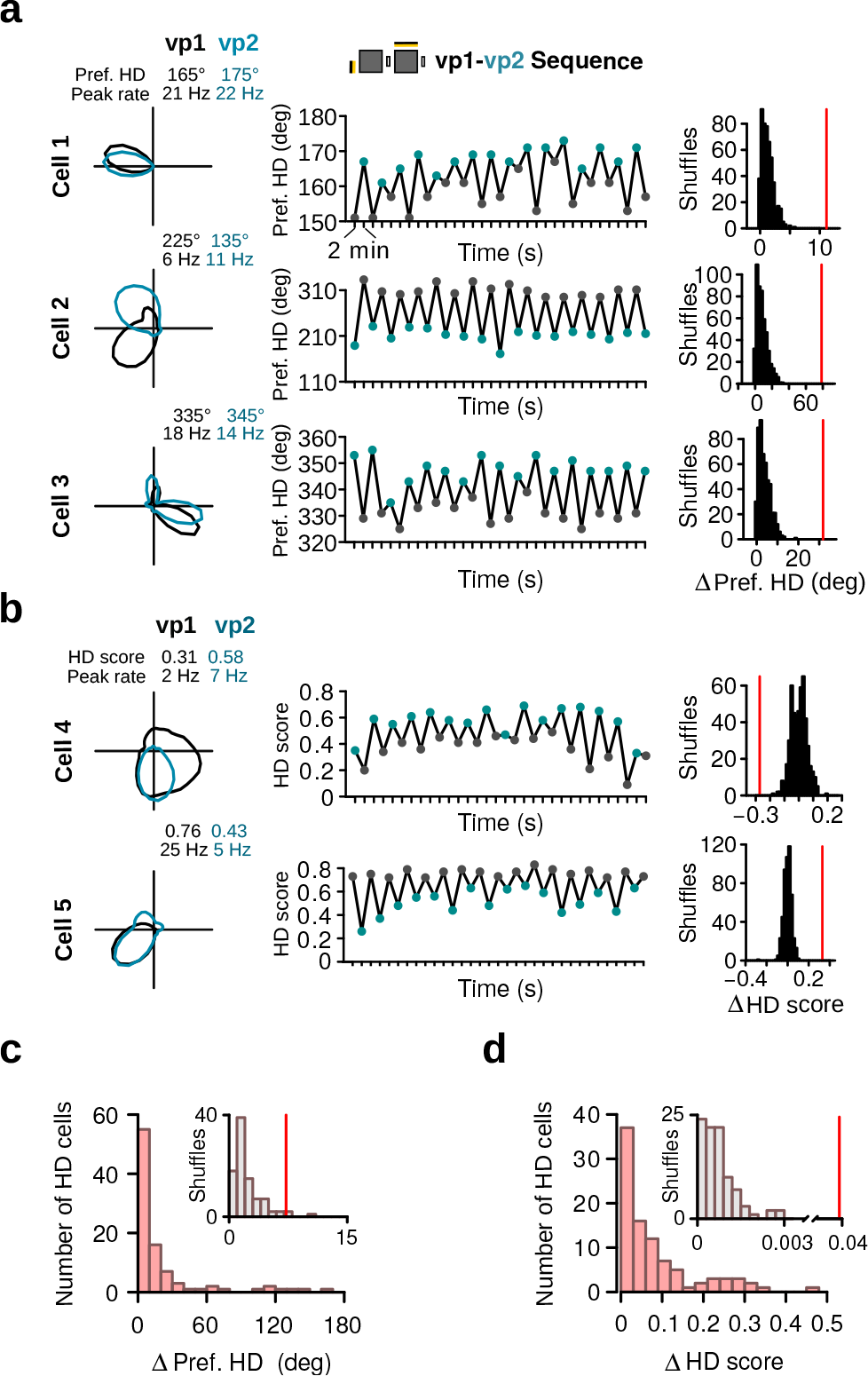
Changes in preferred HD and HD selectivity caused by visual landmarks. **a**, Examples of three HD cells that changed their preferred direction between vp1 and vp2 trials. Left: HD tuning curves during vp1 (black) and vp2 (blue) trials. Numbers indicate peak firing rates and preferred HD during vp1 and vp2 trials. Middle: preferred HD of the same cells during individual 2-min trials. Changes in preferred HD were often readily visible on a single trial basis. Right: observed change in preferred HD between vp1 and vp2 trials (red line) and distribution of preferred HD changes when trial labels were reassigned randomly. **b**, Examples of two HD cells with different HD selectivity during vp1 and vp2 trials. Left: HD tuning curves during vp1 (black) and vp2 (blue) trials. Numbers indicate peak firing rates and HD scores during vp1 and vp2 trials. Middle: HD scores of the same two cells for individual trials. Right: observed difference in HD score between vp1 and vp2 trials (red line) and distribution of HD score differences when trial labels were reassigned randomly. **c**, Distribution of shifts in HD preference between vp1 and vp2 for all HD cells (red). Inset: the median of observed shifts (red line) with the distribution expected by chance (gray). **d**, Same as c but for changes in HD score between vp1 and vp2.

The distribution of changes in preferred direction and HD selectivity across all recorded HD cells was significantly different from chance. This was shown by comparing the observed changes of the HD cells to their respective median changes obtained from the surrogate data. (Figures 2c and 2d; paired Wilcoxon signed-rank test: change in preferred direction, *v* = 127, *P* < 10^−15^; change in HD score, *v* = 4, *P* < 10^−16^).

It was previously shown that HD cells located in the presubiculum and anterodorsal thalamic nucleus maintained their average firing rate when an animal explored different environments (Taube et al., 1990b; Taube and Burton, 1995; Goodridge et al., 1998). We therefore tested whether the mean firing rate of HD cells in the MEC/PaS was also preserved when visual landmarks were manipulated. We observed HD cells that showed pronounced alterations in their firing rate in response to different visual patterns (Figure 3a). For these cells, the instantaneous firing rate oscillated in time, in line with the trial type (Figure 3a). To test whether these changes were significant, we calculated the relative change in firing rate for each neuron: 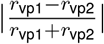. We found that 40.9% (38 out of 93) of the HD cells significantly altered their mean firing rate in response to distinct visual landmarks. At the population level, observed rate changes were larger than those obtained from surrogate data (Figure 3b; paired Wilcoxon signed-rank test, *v* = 12, *P* < 10^−16^). Thus, these results demonstrate that the the mean firing rate of HD cells in the MEC/PaS is modulated by visual landmarks.

**Figure 3.**
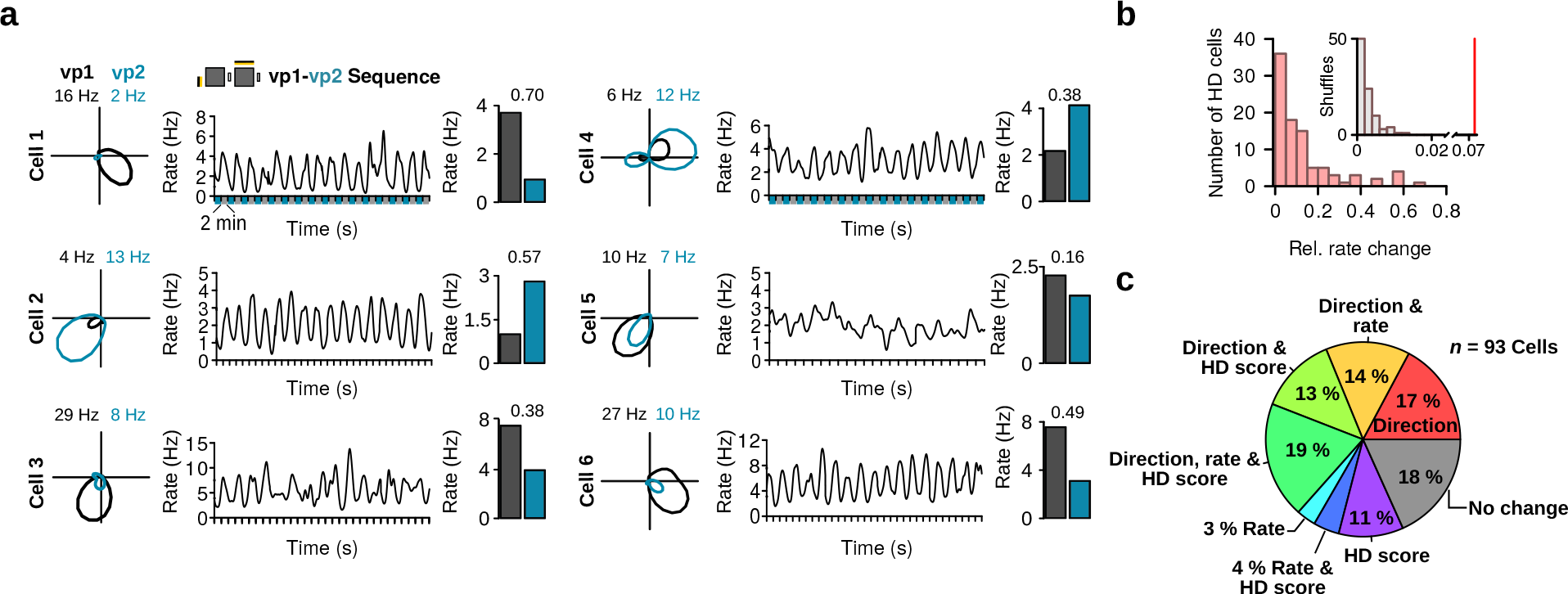
Firing rate changes induced by visual landmarks. **a**, Six HD cells that significantly changed their firing rate depending on visual landmarks. For each cell from left to right: HD tuning curves for vp1 (black) and vp2 (blue) trials, instantaneous firing rates (standard deviation Gaussian kernel 25 s, window size 1 s) as a function of time, and average firing rates during vp1 and vp2 trials. Numbers indicate the relative change in rate of each HD cell. **b**, Distribution of relative change in firing rate for all HD cells (red). Inset: the median of observed rate change (red line) with the distribution expected by chance (gray). **c**, Pie chart illustrating the fractions of HD cells with significant changes in preferred direction, HD score or mean firing rate.

The majority of neurons in the MEC are spatially selective (Diehl et al., 2017). It could therefore be argued that the changes in the firing properties of HD cells between vp1 and vp2 trials were caused by an altered spatial input to HD cells (Cacucci et al., 2004). To rule out this possibility, we focused our analysis on a subset of HD cells with no significant spatial modulation (non-significant sparsity score; 36 out of 93 HD cells). Within this population of non-spatial HD cells, 86.1% (31 out of 36) had a significant change in preferred direction, HD score or mean firing rate between the vp1 and vp2 trials. This proportion was comparable to that observed when considering spatially selective HD cells (Pearson’s Chi-squared test: *χ*^2^ = 0.3543, *df* = 1, *P* = 0.55). Thus, the changes in HD cell properties did not result from an altered spatial signal between vp1 and vp2 trials.

We also tested whether a similar proportion of HD cells showed landmark-driven changes when limiting the analysis to cells recorded from hemispheres in which all tetrodes were positioned in the MEC. We found that 83.6% (56 out of 67 cells) of the MEC HD cells displayed significant changes in preferred direction, HD selectivity or mean firing rate between trial types. This percentage did not significantly differ when analyzing HD cells from hemispheres containing recording sites in the PaS, presubiculum or cortex (76.9%, 20 out of 26 cells, *χ*^2^ = 0.1996, *df* = 1, *P* = 0.66). In summary, the majority of HD cells showed significant changes in preferred direction, HD selectivity or mean firing rate when visual landmarks were manipulated (Figure 3c).

### Bidirectionality of HD cells is affected by visual landmarks

Inspection of the HD cell tuning curves also revealed that a subpopulation of HD cells had two preferred directions (Figure 4a, see also Figures 1e-Cell-4, 2a-Cell-3, 3a-Cell-4). Some of these tuning curves were similar to those previously reported for the retrosplenial cortex (Jacob et al., 2017), but the two peaks were not always at an 180° angle to each other. The bidirectionality of these cells often varied with the trial types (Figure 4a). To quantify these observations, a bidirectionality (BD) score was defined as the firing rate ratio of the two largest peaks in the tuning curve. For some cells, the BD score changed on a trial-by-trial basis (Figure 4a). The bidirectionality in the tuning curves could be observed over short periods of 30 seconds (Figure 4b), suggesting that bidirectionality was not caused by instability of a single preferred direction.

**Figure 4.**
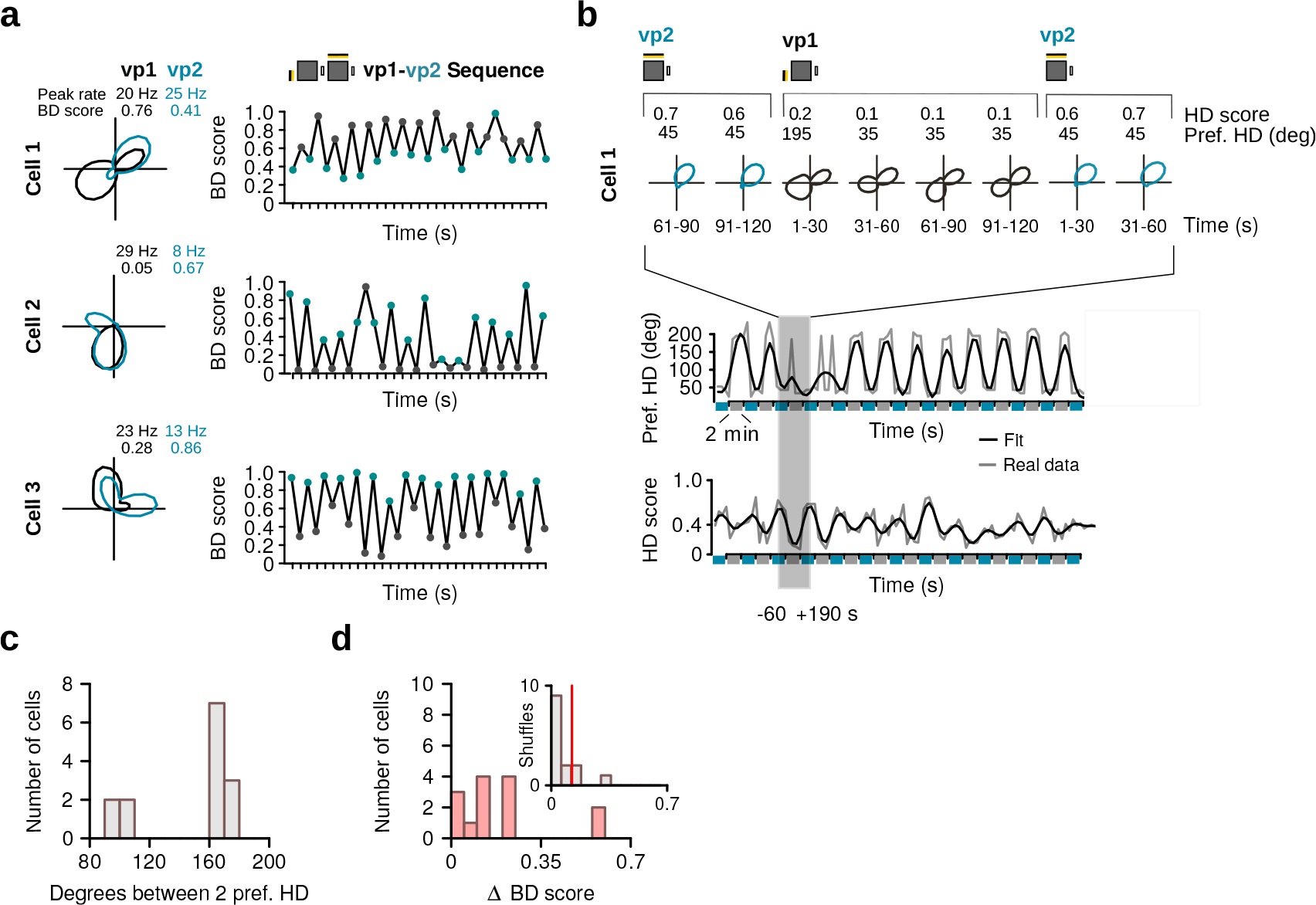
HD cells with bidirectional tuning curves. **a**, Three example HD cells with bidirectional HD tuning curves that changed their bidirectionality with the trial types. From left to right: tuning curves during vp1 and vp2 trials, with numbers indicating the peak firing rates and bidirectionality (BD) scores during vp1 and vp2, and evolution of the BD score during the recording session. Bidirectionality of the tuning curved changed with the two trial types. **b**, Example of a bidirectional HD cell during the transition from vp2 to vp1 trials at a shorter time scale. Top panel: polar plots during 30 second blocks showing that the two preferred HDs are expressed together in short time periods during vp1 trials. Bottom panel: preferred HD and HD score of the cell over time. **c**, Distribution of angles between the two preferred HDs of bidirectional HD cells. **d**, Distribution of change in BD score between vp1 and vp2 for HD cells with a bidirectional HD tuning curve (red, *n* = 14). Inset: the median of observed changes in BD score (red line) with the distribution expected by chance (gray).

Bidirectional HD cells were defined as HD cells for which the BD score was larger than 0.2, the peak firing rates of the two peaks were larger than 2 Hz, and the ratio between the firing rate of the smaller peak and the trough between the peaks exceeded 1.25. These criteria had to be reached during vp1 or vp2 trials. The analysis showed that 15.1% (14 out of 93) of the HD cells were classified as bidirectional, with 2 cells being bidirectional only during vp1 trials, and 6 cells only during vp2 trials. The remaining 6 cells were bidirectional during both trial types. The distribution of angles between the two preferred directions of bidirectional HD cells had two modes close to 90° and 180° degrees (Figure 4c). In some HD cells, the changes in HD score and preferred direction between vp1 and vp2 trials could be explained by a change in bidirectionality (Figure 4a-b). The changes in the BD scores between vp1 and vp2 trials were larger than chance levels (Figure 3d; paired Wilcoxon signed-rank test, *N* = 14, *v* = 12, *P* = 0.009).

### Reorganization of firing associations between HD cells

Computational models of HD cells predict that differences in preferred directions of HD cells are preserved at all times. Support for these models comes from numerous studies showing coherent rotation of the preferred directions of HD cells in response to landmark manipulations (Taube et al., 1990b; Yoganarasimha et al., 2006). However, these findings were obtained from recordings in subcortical areas or in the presubiculum, and it is still not known whether HD cells in the MEC/PaS display the coherence predicted by computational models. We therefore tested whether visually-driven tuning curve changes observed in our protocol were coherent across simultaneously recorded HD cells.

We first inspected the tuning curves of simultaneously recorded HD cells (*N* = 37 HD cell pairs). Figure 5a shows examples of HD cells for which changes from vp1 to vp2 were not coherent. For example, two HD cell pairs modified their difference in preferred direction depending on which visual pattern was presented (HD Pairs 1 and 2). These pairwise changes could be detected during single 2-min trials (Figure 5a, right). A similar uncoupling of simultaneously recorded HD cell pairs was also observed for HD selectivity (Figure 5a, HD Pair 3) and mean firing rate (Figure 5a, HD Pair 4).

**Figure 5.**
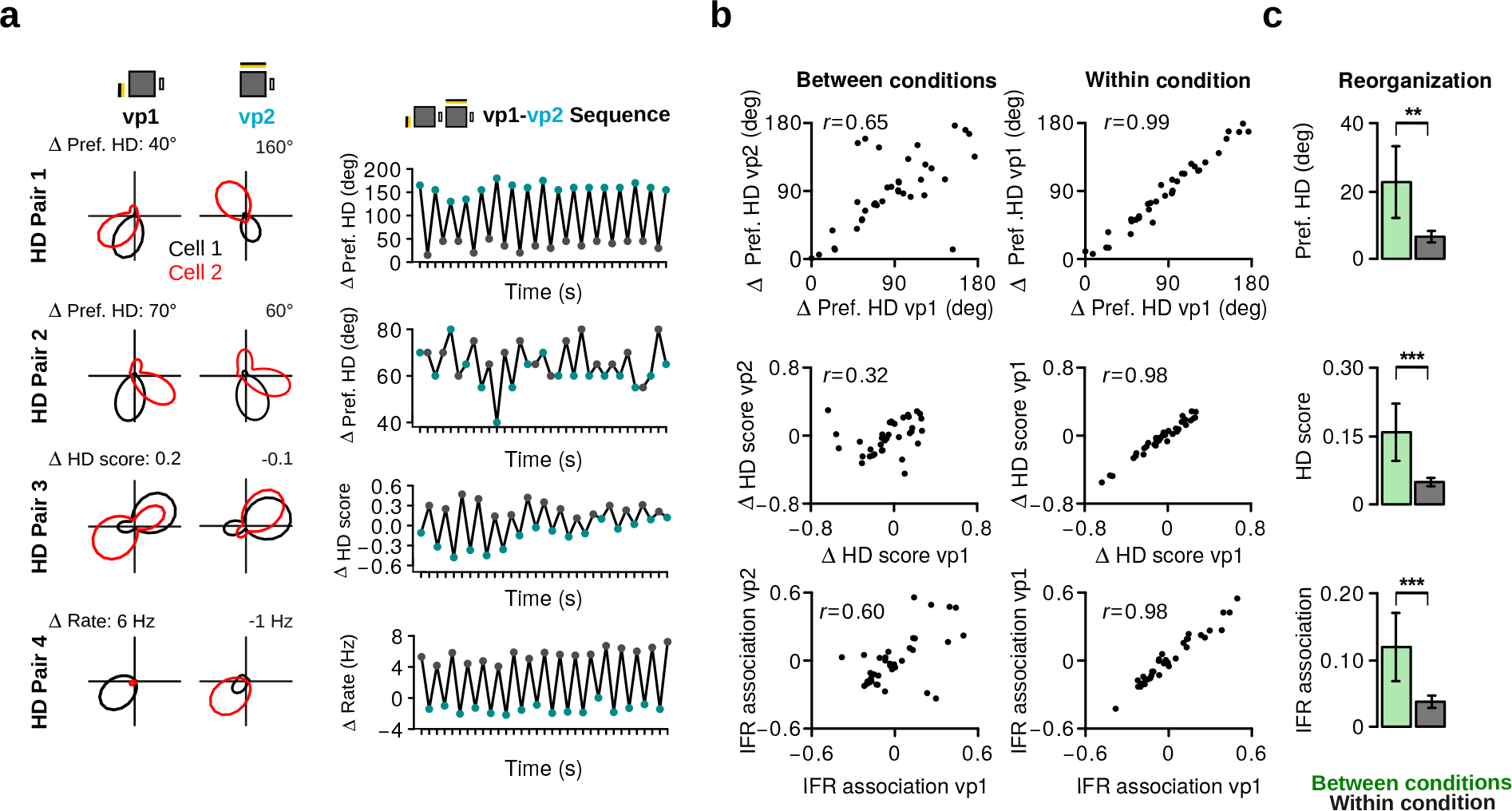
Reorganization of HD cells caused by changes in visual patterns. **a**, Examples of four pairs of HD cells recorded simultaneously showing non-coherent changes. Left: HD polar plots of HD cell pairs during vp1 (first column) and vp2 (second column) trials. The numbers above each polar plot indicate the difference in preferred HD (HD Pairs 1 and 2), in HD score (HD Pair 3) or in mean firing rate (HD Pair 4). Right: temporal evolution of the difference in preferred HD, HD score or mean firing rate for the cell pairs shown on the left. **b**, Correlation between the differences in preferred HD, in HD score and in instantaneous firing rate (IFR) association of HD cell pairs for trials with the same or different visual patterns. Data are shown for vp1 and vp2 trials (left, between conditions) or for two mutually exclusive subsets of vp1 trials (right, within condition). *r* values are correlation coefficients. **c**, Reorganization of preferred HD, HD score and IFR association of HD cell pairs between vp1 and vp2 trials (between conditions) or between two subsets of vp1 trials (within condition). Plots show mean ± 95% confidence intervals. **: *P* < 0.01, ***: *P* < 0.001.

To quantify these non-coherent changes in HD cell activity, we correlated the differences in preferred direction of HD cell pairs observed during vp1 and vp2 trials (between conditions) (Figure 5b, top left panel). As a control, we correlated the same scores obtained from two mutually exclusive subsets of vp1 trials (within conditions) (Figure 5b, top right panel). The correlation coefficient between conditions (*r* = 0.65) was significantly lower than that observed within condition (*r* = 0.99; difference, Fisher Z-transformation: Z-score = 7.72, *P* < 10^−14^). Similarly, differences in HD scores were less correlated between conditions than within conditions (Figure 5b, middle panels; between conditions: *r* = 0.32, within conditions: *r* = 0.98, difference: Z-score = 8.11, *P* < 10^−16^). We also assessed the firing associations of HD cells in the time domain. The instantaneous firing rate (IFR) of each HD cell was calculated and IFR association was defined as the correlation coefficient between IFR vectors of two HD cells. The stability of IFR associations was lower between conditions (*r* = 0.60) than within condition (*r* = 0.98) (Figure 5b, bottom panels; difference: Z-score = 6.62, *P* =10 < 10^−11^).

To further quantify non-coherent changes within the HD cell population, a reorganization score for preferred HD, HD score and IFR association was computed. After obtaining differences in preferred HD, HD score or IFR associations for each cell pair, the absolute difference between the values from vp1 and vp2 trials or from two mutually exclusive subsets of vp1 trials served as reorganization score between or within conditions, respectively (see Materials and Methods). As expected, we observed a larger reorganization between vp1 and vp2 trials than within conditions (Figure 5c, paired Wilcoxon signed-rank test; HD preference: *v* = 146, *P* = 0.00143; HD score: *v* = 115, *P* = 0.00019; firing association: *v* = 135, *P* = 0.00073). These results demonstrate that the internal firing organization of HD cells in the MEC/PaS is modified by visual landmarks. Thus, the activity of HD cells is not entirely constrained by attractor-like dynamics.

### Theta rhythmicity reveals two HD cell populations

A substantial proportion of MEC/PaS neurons show prominent rhythmic activity in the theta frequency band (6-10 Hz) (Cacucci et al., 2004; Mizuseki et al., 2009). This rhythmicity depends on inputs from the medial septum, which are critical for grid cell periodicity in rodents (Mitchell et al., 1982; Brandon et al., 2011; Koenig et al., 2011). We therefore investigated whether HD cells in the MEC/PaS fire rhythmically at theta frequency and whether theta rhythmicity is associated with visually-driven changes in HD response. Theta rhythmicity of each HD cell was quantified from the power spectrum of its instantaneous firing rate (Figure 6a). We found a large variability in the theta rhythmicity of HD cells (Figure 6a). A theta index was calculated by comparing the power in the theta frequency band (6-10 Hz) to that of two adjacent frequency bands (3-5 and 11-13 Hz). The distribution of theta indices for all HD cells is shown in Figure 6b. The theta index distribution was characterized using Gaussian finite mixture models, with the best fitting model having two components (see Materials and Methods; log-likelihood = 98.25, *df* = 5, BIC = 173.83). In line with Alonso and García-Ausst’s work (1987), these two components corresponded to theta rhythmic and non-rhythmic cells (Figure 6b and Supplementary Figure 3a, *N* = 34 non-rhythmic, *N* = 59 theta rhythmic, theta index threshold = 0.07).

**Figure 6.**
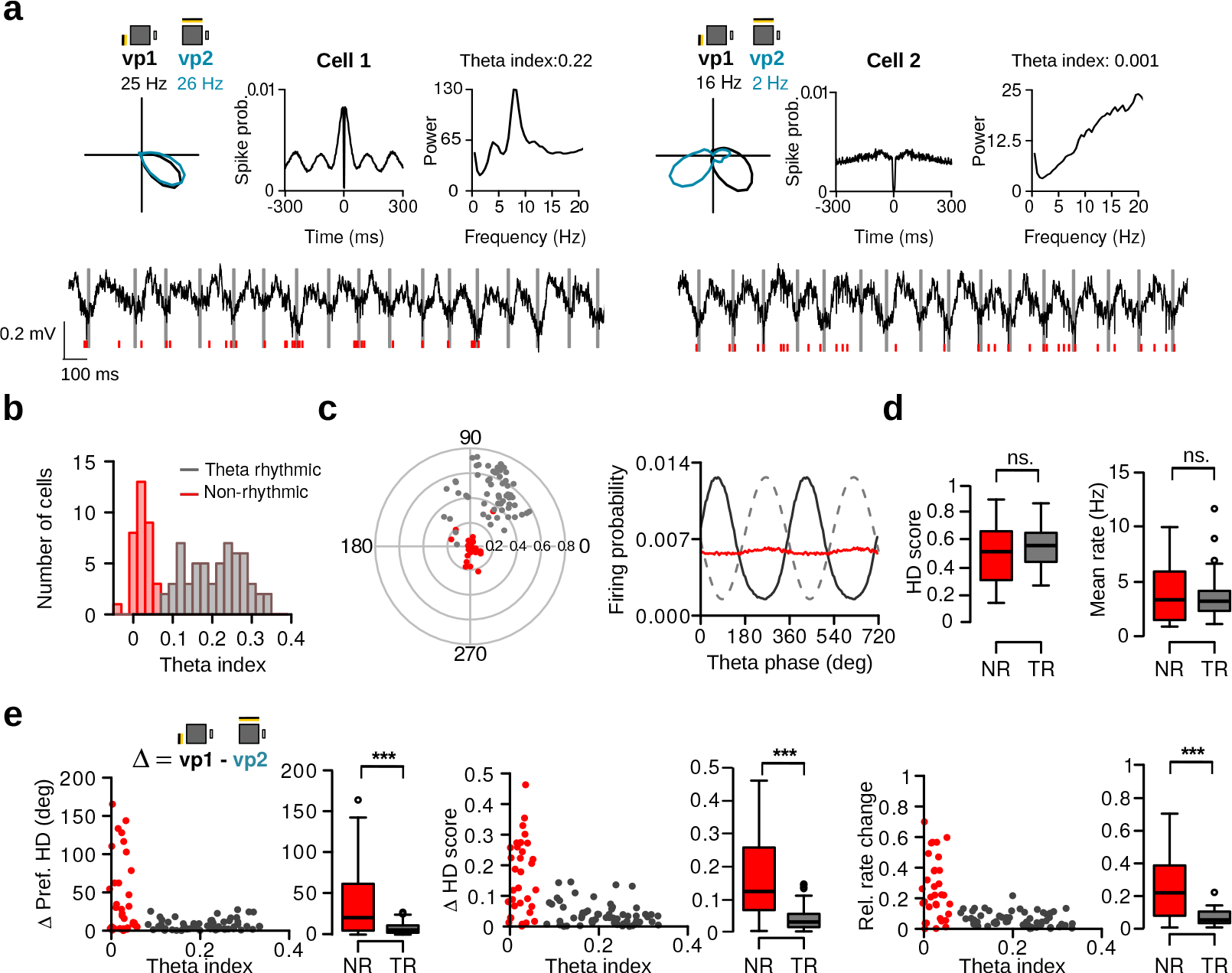
Theta rhythmic activity identifies two populations of HD cells responding differently to visual patterns. **a**, Representative examples of a theta rhythmic HD cell (cell 1) and non-rhythmic HD cell (cell 2). From left to right: HD tuning curves during vp1 and vp2 trials (numbers indicate peak firing rates), spike-time autocorrelations, and power spectra of the instantaneous firing rate. Bottom: raw signals with spike times (red vertical tics) of the HD cells shown above. Vertical gray lines are aligned to the troughs of theta cycles. **b**, Distribution of theta indices of all HD cells. The two populations were identified with a Gaussian finite mixture model. Red: non-rhythmic cells, gray: theta rhythmic cells. **c**), Preferred theta phase and phase locking of HD cell to the LFP theta oscillations. Left: Polar plot showing the preferred theta phase and theta phase modulation of HD cells. Non-rhythmic and theta rhythmic HD cells are depicted in red and gray, respectively. Right: Mean firing probability as a function of theta phase of non-rhythmic (red) and rhythmic (black) HD cells. **d**, HD scores and mean firing rates of non-rhythmic (NR, red) and theta rhythmic (TR, gray) HD cells during vp1 trials. **e**, Scatter plots of the theta index of each HD cell against its change in preferred HD, HD score and relative rate change between vp1 and vp2. Non-rhythmic and theta rhythmic HD cells are depicted in red and gray, respectively. The box-and-whisker plots show the change in preferred HD, HD score and relative rate change between vp1 and vp2 for non-rhythmic and rhythmic HD cells. ns.: not significant, ***: *P* < 0.001.

In addition, the relationship between HD cell firing activity and theta oscillations recorded in the local field potentials was investigated. For each cell, the firing probability as a function of the theta phase, together with the mean vector length and the preferred theta phase of its spikes were calculated. Theta rhythmic HD cells showed much a larger mean vector length than non-rhythmic HD cells (Figure 6c, Wilcoxon rank-sum test, *w* =19, *P* < 10^−15^). Theta rhythmic HD cells fired preferentially at the end of the descending phase of theta oscillations (mean preferred phase: 59.27°; Rayleigh test of uniformity, test statistic = 0.2685, *P* < 10^−19^) whereas non-rhythmic HD cells did not show such preference (Figure 6c, test statistic = 0.2685, *P* = 0.09). The firing rate of all theta rhythmic HD cells was modulated by theta phase. Even though non-rhythmic HD cells displayed weaker theta phase modulation, the firing rate of 94.1% (32 out of 34) of non-rhythmic HD cells was significantly modulated by theta phase.

These large differences in theta modulation were not caused by differences in the local field potential theta power recorded in the proximity of the cell bodies. For each HD cell, we calculated the power spectrum of the local field potentials from its respective tetrode. The power at theta frequency in the local field potentials was similar for theta rhythmic and non-rhythmic HD cells (Supplementary Figure 3b, Wilcoxon rank-sum test, peak theta power: *w* = 1024.5, *P* = 0.87). There was no significant difference between theta indices of cells recorded in hemispheres in which all tetrode tips were located in the MEC compared to those in which some tetrode tips were in the PaS (*w* = 2433.5, *P* = 0.75). The proportion of non-rhythmic to theta rhythmic HD cells also did not change when considering only HD cells recorded simultaneously with a grid cell on the same tetrode (Supplementary Figure 3c, *N* = 15, non-rhythmic: 5 out of 15, 30.0%; theta rhythmic: 10 out of 15, 30.0%; Pearson’s Chi-squared test: χ^2^ < 10^−31^, *df* = 1, *P* = 1.0).

Having established that HD cells of the MEC/PaS can be divided into two classes based on theta rhythmicity, we then compared their HD selectivity and how they responded to changes in visual landmarks. The HD selectivity and mean firing rate of theta rhythmic and non-rhythmic HD cells during vp1 trials were similar (Figure 6d, Wilcoxon rank-sum test, HD score: *w* = 881, *P* = 0.33, mean rate: *w* = 993, *P* = 0.94). Thus, both theta rhythmic and non-rhythmic HD cells displayed similar levels of directional tuning.

We found that the responses of rhythmic and non-rhythmic HD cells to visual landmark manipulations were strikingly different. Non-rhythmic HD cells showed much larger changes in preferred HD between vp1 and vp2 trials than theta rhythmic HD cells (Figure 6e, *w* = 1465, *P* = 0.00012). Likewise, non-rhythmic HD cells also showed larger changes in HD selectivity and mean firing rate between the two trial types than rhythmic HD cells (HD score difference: *w* =1618, *P* < 10^−7^; relative difference in mean firing rate: *w* =1641, *P* < 10^−7^). The percentage of cells with a significant visually-driven change was 74.5% and 94.1% for theta rhythmic and non-rhythimc cells, respectively (Supplementary Figure 3d, χ^2^ = 3.4465, *df* = 1, *P* = 0.06). Taken together, these analyses identified two populations of HD cells that differ in their theta rhythmicity. Moreover, visually-driven changes in tuning curves were much more pronounced in non-rhythmic HD cells compared to theta rhythmic HD cells.

### Non-rhythmic HD cells reorganize their firing associations

The existence of two HD cell populations raises the possibility that HD cells within the theta rhythmic or non-rhythmic population preserved their firing associations when visual landmarks were manipulated. To address this, we compared reorganization scores of HD cell pairs formed by two theta rhythmic or two non-rhythmic HD cells (Figure 7a and 7b). The reorganization scores between conditions (vp1 and vp2 trials) and within condition (vp1 trial subsets) were compared. We found that rhythmic HD cell pairs did not show significant reorganization when visual landmarks were altered (Figure 7b, *N* = 20, preferred HD: *v* = 65, *P* = 0.14, HD score: *v* = 72, *P* = 0.23, IFR association: *v* = 69, *P* = 0.19). In contrast, visual landmark manipulations caused a reorganization of firing associations in non-rhythmic HD cell pairs (*N* = 9, preferred HD: *v* = 2, *P* = 0.01, HD score: *v* = 1, *P* = 0.00781, IFR association: *v* = 0, *P* = 0.00391). These results indicate that the reorganization of firing associations was specific to nonrhythmic HD cells.

**Figure 7.**
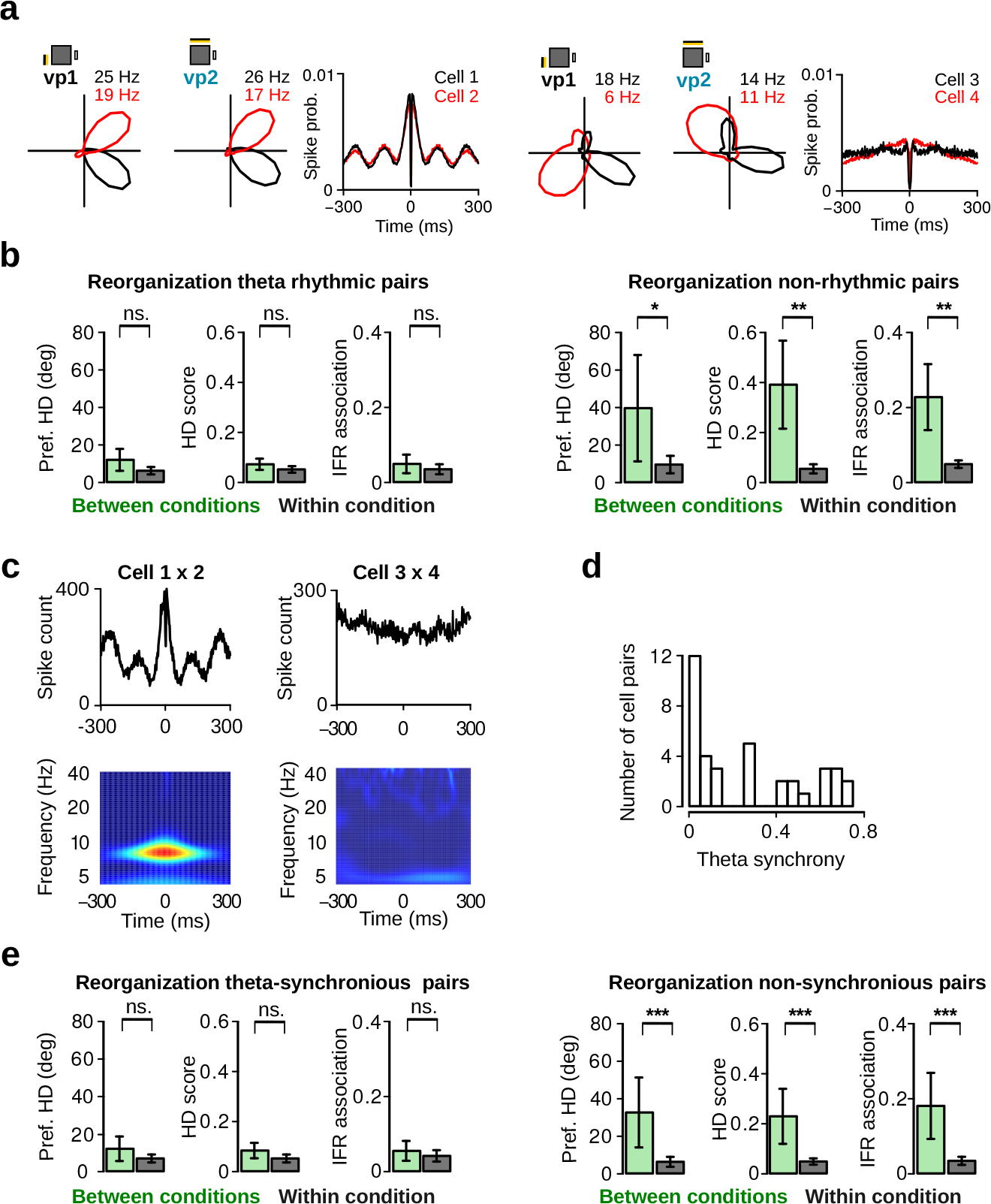
Non-rhythmic HD cells reorganize their firing associations. **a**, Representative example of a theta rhythmic (left) and non-rhythmic (right) HD cell pair. For each pair, from left to right, tuning curves during vp1 and vp2 trials and spike-time autocorrelation of the two neurons are displayed. **b**, Reorganization in preferred direction, HD score and instantaneous firing rate (IFR) association for HD cell pairs formed by two theta rhythmic HD cells (left) or two non-rhythmic cells (right). Reorganization between vp1 and vp2 trials (between conditions) is compared to reorganization between two subsets of vp1 trials (within condition). Plots show mean ± 95% confidence intervals. **c**, Top: spike-time crosscorrelations for the two HD cell pairs shown in A. Below: wavelet transforms of the z-score based spike-time crosscorrelations revealing theta frequency synchronization for the first HD cell pair. Both wavelet transforms were normalized to the same scale from minimal power (dark blue) to maximal power (dark red). **d**, Distribution of theta synchrony scores for all HD cell pairs. **e**, Reorganization for HD cell pairs with high (left) or low (right) theta synchrony. ns.: not significant, *: *P* < 0.05, **: *P* < 0.01,***: *P* < 0.001.

We also tested whether visually-driven reorganization between HD cells can be predicted based on their synchronous activity at theta frequency. We obtained power spectra from spike-time crosscorrelations of simultaneously recorded HD cell pairs (Peyrache et al., 2015) (Figure 7c). Theta synchrony was defined as the average power at theta frequency. From this, the distribution of theta synchrony scores was used to identify theta synchronous and non-synchronous HD cell pairs (Figure 7d, non-synchronous < 0.2, *N* =19, synchronous > 0.2, *N* =18). Theta synchronous HD cell pairs did not show significant visually-driven reorganization (Figure 7e, preferred HD: *v* = 69, *P* = 0.50, HD score: *v* = 51, *P* = 0.14, IFR association: *v* =67, *P* = 0.44). In contrast, HD cell pairs with low co-modulation at theta frequency showed significant reorganization when visual landmarks were altered (preferred HD: *v* =14, *P* = 0.00042, HD score: *v* =16, *P* = 0.00065, IFR association: *v* =18, *P* = 0.00097). These results show that HD cells which synchronized at theta frequency did not undergo visually-driven reorganization. In contrast, non-synchronous HD cells reorganized their firing associations based on visual landmarks.

### Stable firing associations between grid cells

The above results indicate that one population of HD cells in the MEC/PaS show visually-driven reorganization of their firing associations. This contradicts the prediction of current attractor network models of HD cells which states that the firing associations between HD cells are static. Another cell population in this brain region that is thought to be governed by attractor-like dynamics is grid cells (McNaughton et al., 2006). For this reason we sought to determine whether the activity of simultaneously recorded grid cells remained coherent when visual landmarks were manipulated.

We recorded the activity of 219 grid cells during vp1 and vp2 trials (Figure 8a). To investigate whether grid cells altered their firing properties in response to manipulation of visual landmarks, we assessed the stability of their firing rate maps. The firing rate map similarity between the two trial types was significantly lower than the chance level (Figure 8b; paired Wilcoxon signed-rank test, *N* = 219, *v* = 439, *P* < 10^−16^). We found that 48.4% (106 out of 219) of the grid cells showed significant changes in map similarity, preferred HD, or average firing rate between the vp1 and vp2 trials (Figure 8c).

Next, we examined whether the grid cell changes were coherent across simultaneously recorded grid cells (*N* = 223 grid cell pairs, Figure 8d), as predicted by attractor network models. Firing associations based on instantaneous firing rates were highly preserved between vp1 and vp2 trials (Figure 8e, top left panel, *r* = 0.97). Likewise, pairwise map similarity from the vp1 and vp2 trials was also strongly correlated (Figure 8e, bottom left panel, *r* = 0.96). In addition, reorganization levels for pairs of grid cells and pairs of HD cells were compared (Figure 8f). Only HD cells showed significant reorganization of their IFR association and of their firing rate maps between different trial types (reorganization of grid cell pairs between vs. within condition for IFR association: *v* = 10911, *P* = 0.10; map similarity: *v* = 10661, *P* = 0.058). Similar results were obtained when considering only grid cell pairs with at least one grid cells showing significant visually-driven changes of either map similarity, average firing rate or preferred HD (Supplementary Figure 4a-b, *N* = 106 grid cell pairs, IFR association: *v* = 2296, *P* = 0.089; map similarity: *v* = 2247, *P* = 0.064). These findings suggest that the firing associations of grid cells are not significantly altered in response to changes in visual landmarks.

**Figure 8.**
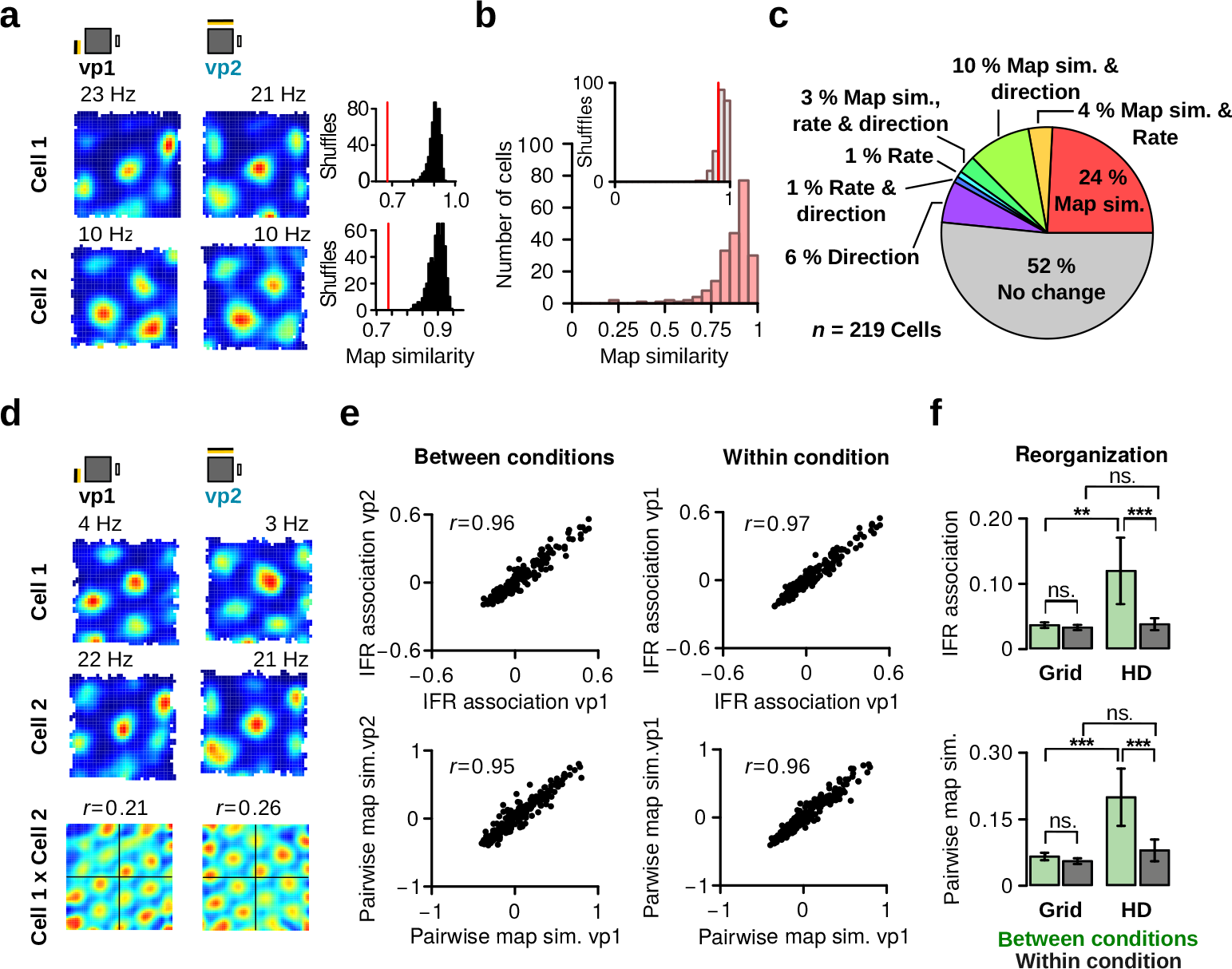
Grid cells retain stable firing associations. **a**, Left: firing rate maps of two representative grid cells during vp1 (left) and vp2 (right) trials. Right: observed map similarity between vp1 and vp2 trials (red line) and distribution of map similarity when trial labels were reassigned randomly. **b**, Distribution of map similarity for all grid cells (red). Inset: the median of observed map similarity (red line) with the distribution expected by chance (gray). **c**, Pie chart illustrating the fractions grid cells with a significant change in map similarity, average firing rate or preferred HD. **d**, Firing rate maps of a two simultaneously recorded grid cells and their spatial crosscorrelation. *r* values are correlation coefficients representing pairwise map similarity of two cells during each trial type. **e**, Comparison of instantaneous firing rate (IFR) associations (top) and pairwise map similarity (bottom) of grid cell pairs during vp1 and vp2 trials (left, between conditions) or for two subsets of vp1 trials (right, within condition). **f**, Reorganization of IFR associations (top) and pairwise map similarity (bottom) between conditions (vp1 vs. vp2) or within condition. Reorganization is shown separately for pairs of grid cells or pairs of HD cells. Plots show mean ± 95% confidence intervals. ns.: not significant, *: *P* < 0.05, **: *P* < 0.01, ***: *P* < 0.001.

## Discussion

HD cells of the MEC/PaS are at the top level of the HD circuit hierarchy. In this brain area, information about direction and running speed is integrated by grid cells to estimate the position of an animal in space (McNaughton et al., 2006; Sargolini et al., 2006; Kropff et al., 2015; Winter et al., 2015). Despite evidence that HD cells are required for grid cell spatial selectivity (Winter et al., 2015), there have been few targeted investigations of the HD signal in the MEC/PaS (Cacucci et al., 2004; Sargolini et al., 2006; Brandon et al., 2013; Giocomo et al., 2014; Tang et al., 2016). Our study identifies several new properties of MEC/PaS HD cells. We found that the majority of HD cells changed their preferred direction, HD selectivity or firing rate in response to distinct visual landmarks. HD cells could be divided into two classes based on whether they fired rhythmically at theta frequency. Non-rhythmic HD cells displayed large and non-coherent responses to visual landmark manipulations. In contrast, theta rhythmic HD cells showed no evidence of landmark-driven reorganization of firing associations.

These properties of MEC/PaS HD cells set them apart from classic HD cells located upstream in the HD system. Indeed, in the anterodorsal thalamic nucleus and presubiculum the peak firing rate and directional tuning of HD cells are unchanged when an animal explored visually distinct environments (Taube et al., 1990b; Yoder et al., 2011). These results suggest that, in these two brain regions, HD cells are driven mainly by self-motion cues derived from vestibular, proprioceptive, and motor efference information (Stackman and Taube, 1997; Yoder and Taube, 2014). The principal effect of visual landmarks on HD cells of the thalamus and presubiculum is to anchor the HD signal to the external world (Skaggs et al., 1995; Yoder and Taube, 2009; Yoder et al., 2011). Our findings demonstrate that the impact of visual information on MEC/PaS HD cells extends beyond this anchoring mechanism. The changes in firing rate of HD cells observed between vp1 and vp2 trials suggests that visual landmarks are an integral part of the signal encoded by some HD cells. We suggest that a significant proportion of HD cells in the MEC/PaS express a conjunctive code for HD and visual landmarks. The origin of this landmark modulation of HD cell activity is still unknown. Two possible origins are other MEC neurons changing their firing patterns depending on visual landmarks (Pérez-Escobar et al., 2016; Diehl et al., 2017), or afferent connections from visual cortical areas (e.g. postrhinal or retrosplenial cortex) (Burwell and Amaral, 1998; Czajkowski et al., 2013; Koganezawa et al., 2015).

A core tenet of HD cell models is that cells within the network have a fixed connectivity and that differences in preferred direction between HD cells do not change (Skaggs et al., 1995; Zhang, 1996; Redish et al., 1996). Although the HD signal is generated at the level of the dorsal tegmental nucleus and the lateral mammillary nucleus, the evidence supporting this assertion has been obtained from recordings performed downstream of these regions (Taube et al., 1990a; Stackman et al., 2003; Yoganarasimha et al., 2006; Peyrache et al., 2015). This suggests that the attractor-like network dynamics of the HD signal generator propagate to downstream areas of the HD system via ascending excitatory connections (Peyrache et al., 2015). In the current study, we found that the firing pattern of HD cells within the MEC/PaS contradicted this view. Indeed, the firing associations of non-rhythmic HD cells reorganized when visual landmarks were changed. In contrast, no evidence of reorganization was observed in theta rhythmic HD cells under similar conditions. Our data point to the existence of two distinct HD cell populations within the MEC/PaS that hold different reorganization capabilities. Interestingly, a recent study also reported the existence of at least two populations of HD cells in the dysgranular retrosplenial cortex, and one of which expressed bidirectional tuning curves that were controlled by local landmarks (Jacob et al., 2017). Taken together with our findings, this indicates that the HD signal in cortical areas is more heterogeneous than in the anterodorsal thalamus or the presubiculum.

The different firing properties of theta rhythmic and non-rhythmic HD cells suggest that they support different functions. Theta rhythmic HD cells showed no reorganization during our experiment. This coherent HD signal could be well suited for providing directional information to grid cells. Indeed, computational models of grid cells assume that the HD signal remains coherent, and recordings from grid cells indicates that they do not show major reorganization after visual cue manipulations (Hafting et al., 2005; Pérez-Escobar et al., 2016). The theta modulation of rhythmic HD cells might also play an important role in the integrations of the HD signal by grid cells. It has been proposed that the HD signal needs to be transformed into a theta-modulated signal to be used for navigation (Cacucci et al., 2004). Moreover, the segregation of HD cell ensembles during individual theta cycles might also contribute to grid periodicity (Brandon et al., 2013). We therefore suggest that an important function of theta rhythmic HD cells is to provide a directional input to the grid cell network.

One function of non-rhythmic HD cells might be to act as an interface between visual landmarks and the coherent HD signal. These non-rhythmic HD cells show large, non-coherent tuning changes upon landmark manipulations. An interesting mechanism that links visual landmarks to the HD signal has been put forward by Jacob and colleagues based on their recordings in the dysgranular retrosplenial cortex (Jacob et al., 2017). They suggest that landmark-controlled HD cells could mediate a two way interaction between visual landmarks and a coherent HD signal. This idea can be applied to the two HD cell populations identified in our study. The tuning curves of non-rhythmic HD cells would then be established partly based on landmark-dependent inputs. The HD selectivity of these cells would make them co-active with a subgroup of classic HD cells (e.g. theta rhythmic HD cells) with similar HD preferences. Over time, if the visual landmarks controlling the non-rhythmic HD cells remain stable, the association between the non-rhythmic HD cells and the coherent HD signal would be strengthened, allowing stable visual landmarks to gain preferential control over the coherent HD signal. This way, the population activity of landmark-controlled HD cells could contribute to setting the preferred directions of theta rhythmic HD cells when an animal enters visually distinct environments. One interesting possibility is that this interaction between non-rhythmic and rhythmic HD cells might contribute to the orientation of grid cells within each grid module (Stensola et al., 2012).

In conclusion, our results establish that HD cells of the MEC/PaS comprise two distinct types of cells. Theta rhythmic HD cells react coherently when visual landmarks are altered and resemble classic HD cells located upstream in the HD system. In contrast, non-rhythmic HD cells not only show larger responses to visual landmark manipulations, but also respond non-coherently to these manipulations. This study therefore reveals the existence of landmark-driven reorganization within the HD system in the parahippocampal region. Previous work has shown that non-periodic place selective neurons of the MEC react to alterations in the recording environment by shifting the location of their firing fields (Diehl et al., 2017). The findings presented here indicate that a similar context-dependent reorganization exists within the HD code.

## Materials and Methods

The raw data (spike trains, position data and histology) together with the computer code of this study will be freely available on digital repositories upon publication.

### Subjects

The subjects were nine 3-6 month-old male wild type C57BL/6 mice. They were singly housed in 26 cm × 20 cm × 14 cm high cages containing 2 cm of saw dust and one facial tissue. Mice were kept on a 12-h light-dark schedule with all procedures performed during the light phase. All experiments were carried out in accordance with the European Committees Directive (86/609/EEC) and were approved by the Governmental Supervisory Panel on Animal Experiments of Baden Wurttemberg in Karlsruhe (35-9185.81/G-50/14).

### Surgical procedure

Mice were implanted with two 4-tetrode microdrives, with one targeted at the MEC/PaS of each hemisphere. The microdrives allowed independent movement of individual tetrodes, which were made from four 12-*μ*m tungsten wires (California Fine Wire Company). Mice were anaesthetized with isoflurane (1-3%) and fixed to the stereotaxic instrument. The skull was exposed and 4 miniature screws were inserted into the skull. Two screws located above the cerebellum served as ground electrodes. The skull above the MEC/PaS was removed and the tetrode bundles were implanted at the following coordinates (ML: ±3.1 mm from the midline, AP: 0.2 mm anterior from the transverse sinus, 6° angle in the posterior direction). Once the tetrode tips were 0.8 mm into the cortex, the microdrives were fixed to the skull with dental cement. During the first 72 hours post surgery, mice received a s.c. injection of temgesic (0.1 mg/kg) every 8 hours. Mice were given a week to recover after surgery.

### Recording system, spike extraction and spike clustering

Mice were connected to the data acquisition system (RHD2000-Series Amplifier Evaluation System, Intan Technologies, analog bandwidth 0.09-7603.77 Hz, sampling rate 20 kHz) via a lightweight cable. Electrode signals were amplified and digitized by two 16-channel amplifier boards (RHD2132, Intan Technologies). Recording was controlled using ktan software (https://github.com/kevin-allen/ktan). Action potential detection was performed offline from the bandpass-filtered signal (800-5000Hz). Waveform parameters were extracted with principal component analysis. Clusters of spikes were automatically generated using Klustakwik (https://sourceforge.net/projects/klustakwik/), before being manually refined with a graphical interface program.

Clusters quality was estimated from the spike-time autocorrelation and isolation distance. A refractory period ratio was calculated from the spike-time autocorrelation from 0 to 25 ms (bin size: 0.5). The mean number of spikes from 0 to 1.5 ms was divided by the maximum number of spikes in any bin between 5 and 25 ms. Any cluster with a refractory period ratio larger than 0.125 was discarded. Likewise, clusters with an isolation distance smaller than 5 were discarded.

The position and HD of the mouse was estimated from the position of two infrared-LEDs (wave length 940 nm), one large and one small, that were attached to one amplifier. The large and small LEDs were located anterior and posterior to the center of the head, respectively. The distance between the LEDs was approximately 8 cm. An infrared video camera (resolution of 10 pixels/cm, DMK 23FM021, The Imaging Source) located directly above the recording environment monitored the LEDs at 50 Hz. The location and HD of the mouse were extracted on-line from the position of the LEDs (https://github.com/kevin-allen/positrack).

### Initial training

Initial training began one week after surgery and took place in a different room than that used for recording sessions. Food intake was controlled to reduce the weight of the mice to 85% of their normal free-feeding weight. Mice were trained to retrieve food rewards (AIN-76A Rodent tablets 5 mg, TestDiet) delivered at random location within a 70 × 70 cm open field, 3 times per day for 10 min. After 2 days of training, the training time was extended (3 × 15min) and the microdrives were connected to the recording system. This procedure continued until the mice explored the entire open field within 15 min. On each day, the raw signals were monitored on an oscilloscope and tetrodes were lowered until large theta oscillations were visible on most tetrodes.

### Recording environment

The recording environment consisted of an elevated gray square platform (70 × 70 cm) made of PVC. The platform was surrounded by four 40-cm-high walls located 10 cm away from the edges of the platform. A standard white cue card (21 × 29.7 cm) was attached to the center of one wall and remained in place throughout the experiment. Two visual patterns made of LED strips (color temperature: 3000K, Ribbon Slim Top, Ledxon Group, powered by six 1.2 V batteries) were affixed to two other adjacent walls. Visual pattern 1 (vp1) consisted of 4 horizontal 20-cm-long rows (5 cm between rows) starting 7 cm away from the junction of two walls. Visual pattern 2 (vp2) was a single horizontal 80-cm-long row of 48 LEDs. These two visual patterns were the only sources of visible light in the recording environment, and were switched on and off via a relay switch operated by a microcontroller (Arduino Uno).

The recording environment was surrounded by opaque black curtains. Food rewards (AIN-76A Rodent tablets 5 mg, TestDiet) were delivered at random locations from a pellet dispenser located above the ceiling of the recording environment (CT-ENV-203-5 pellet dispenser, MedAssociates). The pellet dispenser was controlled by a microcontroller (Arduino Uno) and the inter-delivery intervals ranged from 20 to 40 s.

### Recording protocol

The mouse was brought to the recording room at the beginning of the recording session and vp1 was turned on. The mouse was placed on the square platform and allowed to forage for 20 min. The presented visual pattern alternated between vp1 and vp2 every 2 min. After 20 min, the mouse was taken off of the platform and placed in a small rest box (23 × 25 × 30 cm) for 20 min. No food reward was available in the rest box. The rest of the recording session comprised a series of 20 min blocks with the following sequence: foraging-rest-foraging-rest-foraging. In each recording session, the mouse foraged on the square platform for a total of 80 min. At the end of the recording session, the tetrodes were lowered by 25-50 *μ*m.

### Identification of spatially selective neurons

Most functional cell types were identified from their HD tuning curves or firing rate maps. A HD tuning curve consisted of the firing rate of a cell as a function of HD (10° bins). An occupancy vector containing the time in seconds spent within the different HD bins was calculated and smoothed with a Gaussian kernel (standard deviation of 10°). The number of spikes falling within the same bins were then divided by the occupancy vector to obtain the firing rate for different HDs. The firing rate vectors were smoothed with a Gaussian kernel (standard deviation of 10°).

Firing rate maps were generated by dividing the square platform in 2 × 2 cm bins. The time in seconds spend in each bin was calculated and this occupancy map was smoothed with a Gaussian kernel (standard deviation of 3 cm). The number of spikes in each bin was divided by the smoothed occupancy map to obtain a firing rate map. A smoothing kernel (standard deviation of 3 cm) was applied to the firing rate map. Only periods when the mouse ran faster than 3 cm/s were considered.

To identify HD cells, vp1 and vp2 tuning curves were calculated by concatenating data from all vp1 and all vp2 trials, respectively. A HD score was defined as the mean vector length of the tuning curve. The preferred direction of the neuron was the circular mean of the tuning curve. HD cells had to have a HD score larger than 0.4 and a peak firing rate larger than 5 Hz during vp1 trials or vp2 trials or both to be selected. The arbitrary threshold of 0.4 is more conservative than most thresholds obtained with a shuffling procedure. The arbitrary threshold was used to ensure that neurons classified as HD cells had strong HD selectivity and clear preferred HDs.

A third criterion was added to the definition of HD cells to ensure that their HD tuning curve was not simply a consequence of spatial selectivity coupled with a biased sampling of HD on the maze. This was required because the behavior of mice is not completely random and not all HDs are equally sampled at every position on the platform. We therefore applied the distributive hypothesis method (Muller et al., 1994; Cacucci et al., 2004) which measured the similarity between the observed HD tuning curve and an expected tuning curve derived from the firing rate map of the cell and the HD occupancy probability at every bin of the firing rate map. The predicted HD tuning curve was defined as follows:

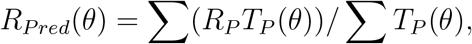

where *R_P_* was the firing rate in one bin of the firing rate map and *T*_*p*_(*θ*) was the time spent facing HD *θ* in that bin. A distributive ratio, *DR*, was then calculated to estimate the similarity between the observed and predicted HD tuning curve:

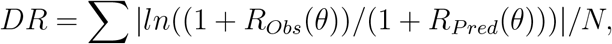

where *N* was the number of bins in a HD tuning curve. A DR of zero indicated that the observed HD tuning curve was well predicted by a combination of spatial selectivity and bias in HD sampling. Higher value indicated that the HD tuning curve was poorly predicted by the spatial selectivity of the cell and that its firing rate was modulated by HD. Cells had to have a DR larger than 0.2 to be considered putative HD cells.

Grid cells were identified based on the periodicity in each firing rate map. A spatial autocorrelation matrix was calculated from the firing rate map. Peaks in the autocorrelation matrix were defined as more than 10 adjacent bins with values larger than 0.1. The 60° periodicity in the spatial autocorrelation matrix was estimated using a circular region of the spatial autocorrelation matrix containing up to six peaks and excluding the central peak. Pearson correlation coefficients (*r*) were calculated between this circular region and a rotated version of the same region (by 30°, 60°, 90°, 120°, and 150°). A grid score was obtained from the formula:

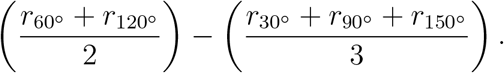

Significance thresholds for grid scores were obtained by shifting the position data by at least 20 s before recalculating grid scores. This procedure was repeated 100 times for each neuron to obtain surrogate distributions. The 99th percentiles of the null distributions were used as significance thresholds.

Spatial selectivity was measured using a sparsity score, adapted so that high scores reflected high sparsity:

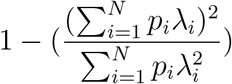

where *N* was the number of bins in the firing rate map, *p_i_* and λ_*i*_ were the occupancy probability and firing rate in bin *i*, respectively. Significance levels for sparsity score were obtained with the same shuffling procedure as for grid scores.

The firing rate modulation by running speed was estimated using the speed score (Kropff et al., 2015), which was defined as the correlation coefficient between the running speed of the animal and the instantaneous firing rate of a neuron. The instantaneous firing rate was calculated by counting the number of spikes observed in 1 ms time bins before applying a Gaussian smoothing kernel (standard deviation of 100 ms).

The firing probabilities were integrated over 100 ms time bins and transformed to firing rates. The running speed of the animal was obtained for these 100 ms time bins and a Pearson correlation was performed between the running speed and the instantaneous firing rate. Only time windows in which the running speed was between 3 and 100 cm/s were considered. Significance levels for speed scores were obtained with the same shuffling procedure as for the grid scores.

### Statistical significance of visually-driven changes

To determine if a HD cell changed significantly their preferred HD, HD score, or mean firing rate between vp1 and vp2 trials, a shuffling procedure was used. The trial labels vp1 and vp2 were reassigned randomly to the forty 2-min trials before recalculating vp1 and vp2 tuning curves. The difference in preferred HD, HD score and mean firing rate between the reassigned vp1 and vp2 trials were calculated. These last steps were repeated 500 times to obtain a distribution of changes expected to occur by chance for the 3 variables. The observed changes of the HD cell were considered significant if they were larger than 99% of the differences obtained with the shuffling procedure. Because the firing properties of individual cells influenced the shuffled distributions, the changes observed in one cell were only compared to the shuffled data of that same cell.

### Bidirectionality of HD tuning curves

Bidirectionality in HD tuning curves was determined from a bidirectionality (BD) score. BD score was defined as the firing rate ratio between the two largest peaks in the tuning curve. To be considered a peak, the firing rate had to be above 2 Hz. Bidirectional HD cells were identified as HD cells for which the BD score was larger than 0.2 and the ratio between the firing rate of the smaller peak and the trough between the peaks exceeded 1.25.

### Instantaneous firing rate for firing associations

The instantaneous firing rate (IFR) of a cell was obtained by counting the number of spikes in 1 ms time bins and applying a Gaussian smoothing kernel (standard deviation of 200 ms) to this spike vector. The firing probability was integrated over 100 ms time windows and transformed to a firing rate. The IFR association of two neurons was estimated by the correlation coefficient between their respective IFR vectors.

### Cell pair reorganization between trial types

Reorganization scores were defined to investigate whether the firing associations between pairs of cells changed between vp1 and vp2 trials. For a cell pair, cell *i* and *j*, the preferred HD (*PrefHD*) reorganization score between conditions was defined as follows:

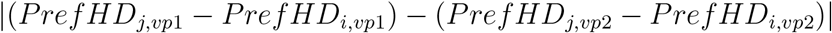

As a control, a reorganization score was calculated using two mutually exclusive subsets of vp1 trials (*vp*1.l and *vp*1.2):

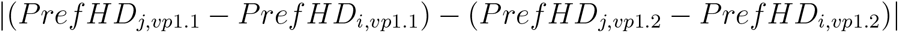

A similar procedure was used to calculate reorganization of HD scores by replacing the preferred HD by HD score. Reorganization of IFR associations was defined as the absolute difference between IFR associations observed during vp1 and vp2 trials.

### Theta rhythmicity and theta oscillations

The theta rhythmicity of neurons was estimated from the instantaneous firing rate of the cell. The number of spikes observed in 1 ms time window was calculated and convolved with a Gaussian kernel (standard deviation of 5 ms). The firing probability was integrated over 2 ms windows and transformed into a firing rate. A power spectrum of the instantaneous firing rate was calculated using the *pwelch* function of the oce R package. The estimates of the spectral density were scaled by multiplying them by the corresponding frequencies: *spec*(*x*) * *freq*(*x*). A theta rhythmicity index for each neuron was defined as 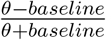, where *θ* is the mean power at 6-10 Hz and *baseline* is the mean power in two adjacent frequency bands (3-5 and 11-13 Hz). The theta rhythmicity indices of HD cells were analyzed with the *Mclust* function of the R package *mclust* which uses Gaussian mixture modeling and the EM algorithm to estimate the number of components in the data.

Theta oscillations were detected using one wire of every tetrode. The signal was bandpass filtered at delta (2-4 Hz) and theta (6-10 Hz) frequencies and the power of the filtered signals (root mean square) was calculated in non-overlapping 500 ms windows. Windows with a theta/delta power ratio larger than 2 were defined as theta epochs. Individual theta cycles within theta epochs were identified using the bandpass filtered (5-14 Hz) signal. The positive-to-negative zero-crossing in the filtered signal delimited individual theta cycles and were assigned phases 0 and 360. The theta phase of spikes was linearly interpolated within the cycle boundaries. For each neuron, the modulation of firing rate by theta oscillations was quantified by calculating the mean vector length of the theta phases assigned to the spikes. The circular mean of the spikes served as the preferred theta phase of a neuron.

Co-modulation of spike activity by theta oscillations for pairs of HD cells was estimated from their spike-time crosscorrelation (-300 to 300 ms, bin size: 2 ms). Each crosscorrelation was transformed into a vector of Z-scores. A discrete wavelet transform (*WaveletComp* R package) was applied to Z-score vector. The mean power in the theta frequency band (6 to 10 Hz) served as theta synchronization score.

### Histology

At the end of the experiment, mice were deeply anesthetized with an i.p. injection of ketamine/xylazine, and perfused with saline followed by 4% paraformaldehyde. The brains were removed and stored at 4° C in 4% paraformaldehyde for at least 24 hours before being sectioned in 50 *μ*m thick slices and stained with cresyl violet. All brain sections were digitized with a motorized widefield slide scanner (Axio Scan.Z1, Zeiss).

## Acknowledgements

We thank H. Monyer, M.I. Schlesiger, and D.A.A. MacLaren for their constructive comments on the manuscript, and J.A. Pérez-Escobar, J. Ivanova and Y. Wang for their technical assistance. This work was supported by an Emmy Noether Program grant (AL 1730/1-1) and a Collaborative Research Centre (SFB-1134) from the Deutsche Forschungsgemeinschaft.

## Author contributions

O.K. and K.A. designed the study; O.K., P.L., L.K. and K.A. performed surgeries and recordings; O.K. performed data analysis; O.K. and K.A. wrote the manuscript in consultation with all authors; K.A. provided supervision and obtained funding.

## Supplementary Information

**Supplementary Figure 1.**
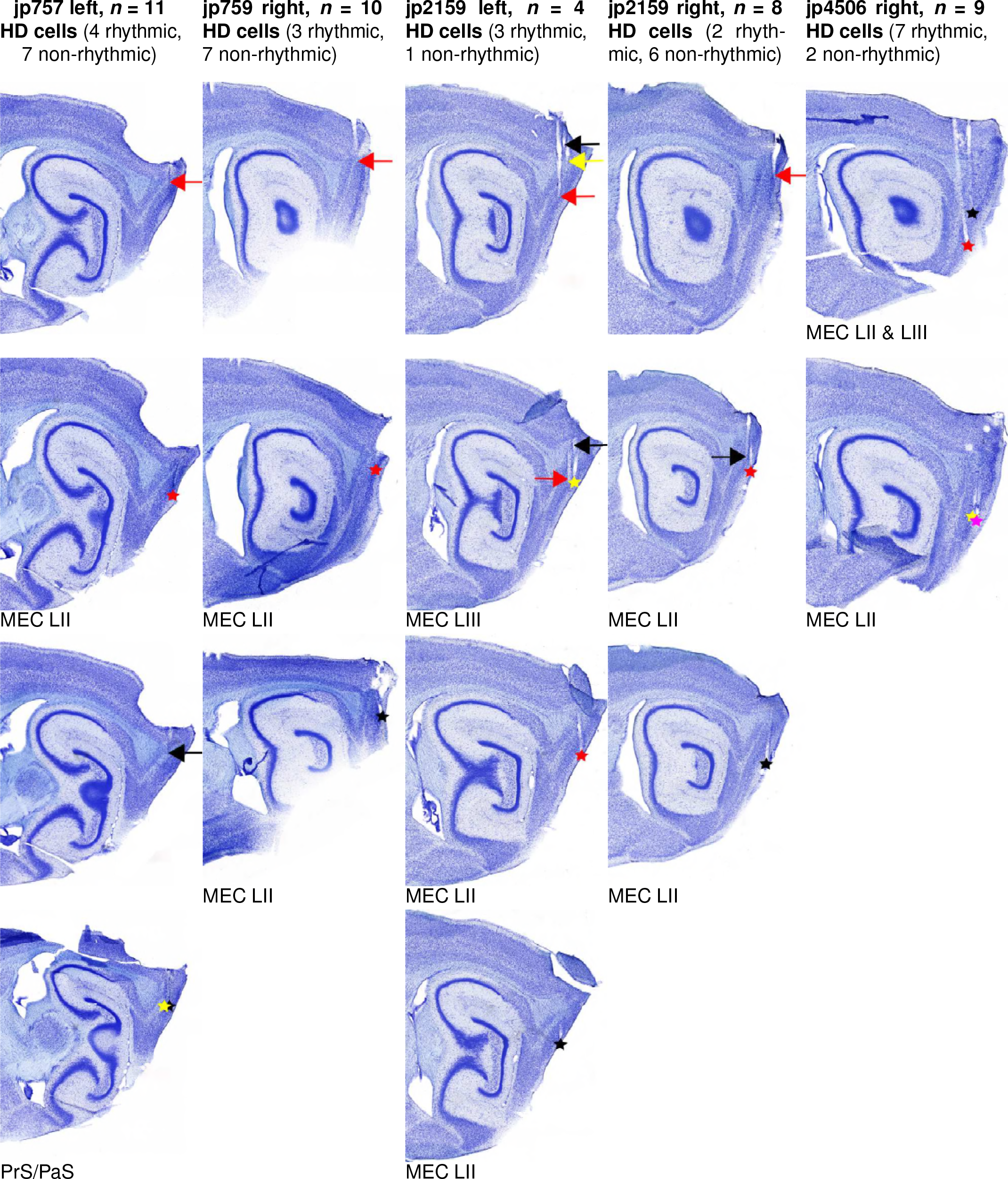
Examples of recording sites. Each column shows sagittal brain sections from one hemisphere. Arrows point to the tetrode tracks and asterisks show the location of the tetrode tips at the end of the experiment. For each hemisphere, a different color was assigned to each tetrode. The brain regions in which the tetrode tips were found are indicated below each section.

**Supplementary Figure 2.**
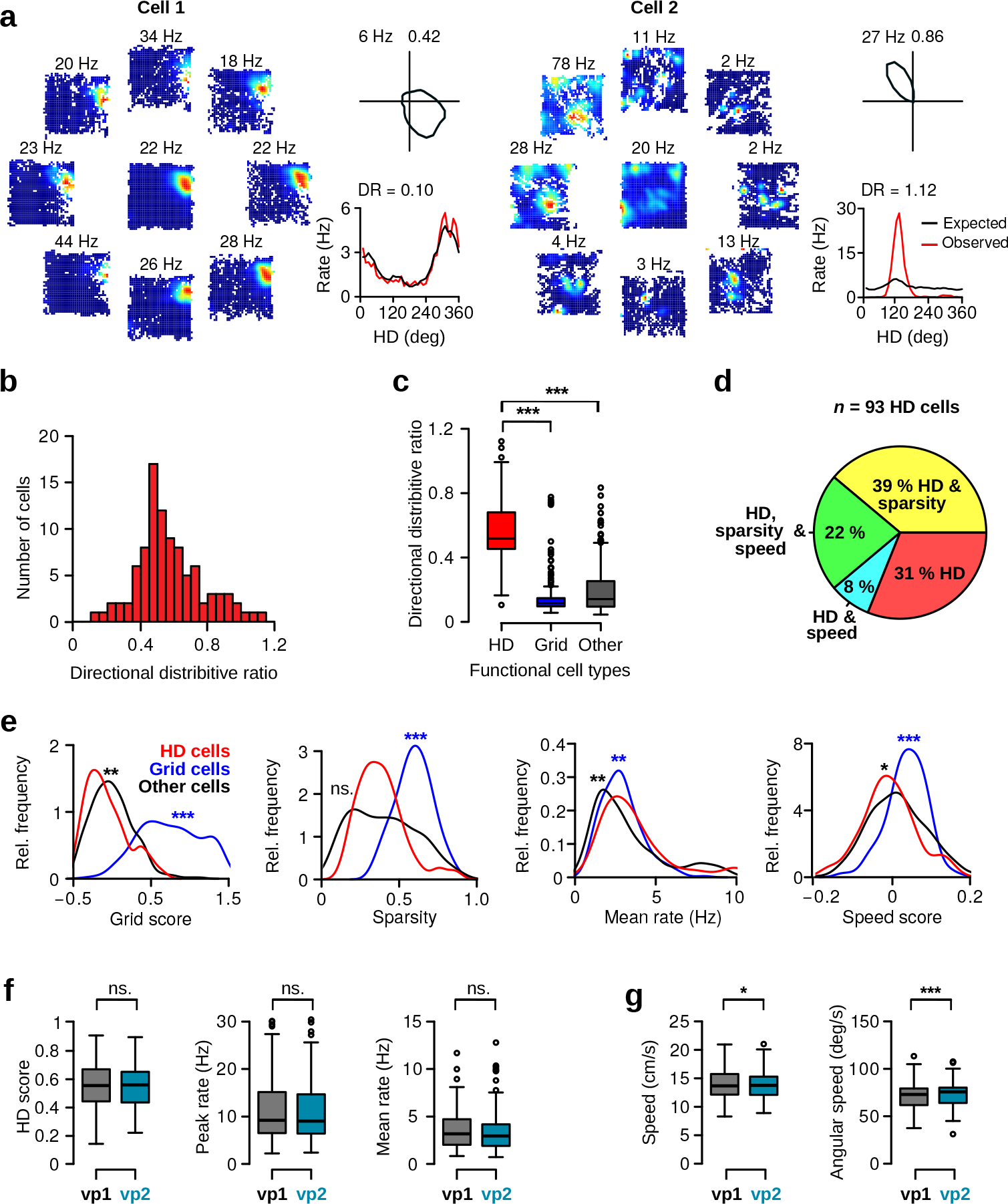
Directional distributive ratio, HD cell properties and behavior during vp1 and vp2 trials. **a**, Directional distributive ratio (DR) for two representative HD cells with low (cell 1) and high DR (cell 2). Left: HD dependent spatial firing rate maps. The central firing rate map is direction independent. Surrounding maps show firing rate maps for specific HDs in 45 deg bins. Top right: polar plot showing the HD tuning curve of the neuron. Numbers indicate peak firing rates and HD scores. Bottom right: Observed (red lines), expected firing rate (black lines) as function of HD and corresponding DR scores. A DR near 0 indicates that the observed tuning curve of a neuron can be explained by its spatial selectivity. **b**, Distribution of DR for all putative HD cells. **c**, Box-and-whisker plots showing DR across different functional cell types: HD cells (HD), grid cells (Grid, *n* = 219), and other low firing rate cells (other, firing rate < 10 Hz, *n* = 335). DR was computed during vp1 trials. HD cells had significantly higher DR compared to other functional cell types (Wilcoxon rank-sum test, HD vs. grid cells, *W* = 20085, *P* < 10^−16^, HD vs. other cells *W* = 29717, *P* < 10^−16^). **d**, Pie chart illustrates the fractions of HD cells with significant spatial sparsity or speed scores. **e**, Distributions of grid scores, sparsity scores, mean firing rates and speed scores for HD cells (red lines), grid cells (blue lines), and other cells (black lines). Asterisks indicate significant difference in scores between HD and grid cells (blue), and HD and other cells (black). **f**, HD scores, peak and mean firing rates for HD cells (paired Wilcoxon signed-rank test, *n* = 93, HD score: *V* = 2015, *P* = 0.52; peak firing rate: *V* = 2267, *P* = 0.76; mean firing rate: *V* = 2234, *P* = 0.85). **g**, Average running speed (left) and average head angular velocity (right). Magnitude of changes between the two trial types was small (paired Wilcoxon signed-rank test, *n* = 68 recording sessions containing HD cells, median running speed, vp1: 13.7 cm/s, vp2: 13.8 cm/s, *V* = 788, *P* = 0.02; median head angular speed, vp1: 73.0 deg/s, vp2: 75.7 deg/s, *V* = 522, *P* < 10^−5^). ns.: not significant, *: *P* < 0.05, **: *P* < 0.01, ***: *P* < 0.001.

**Supplementary Figure 3.**
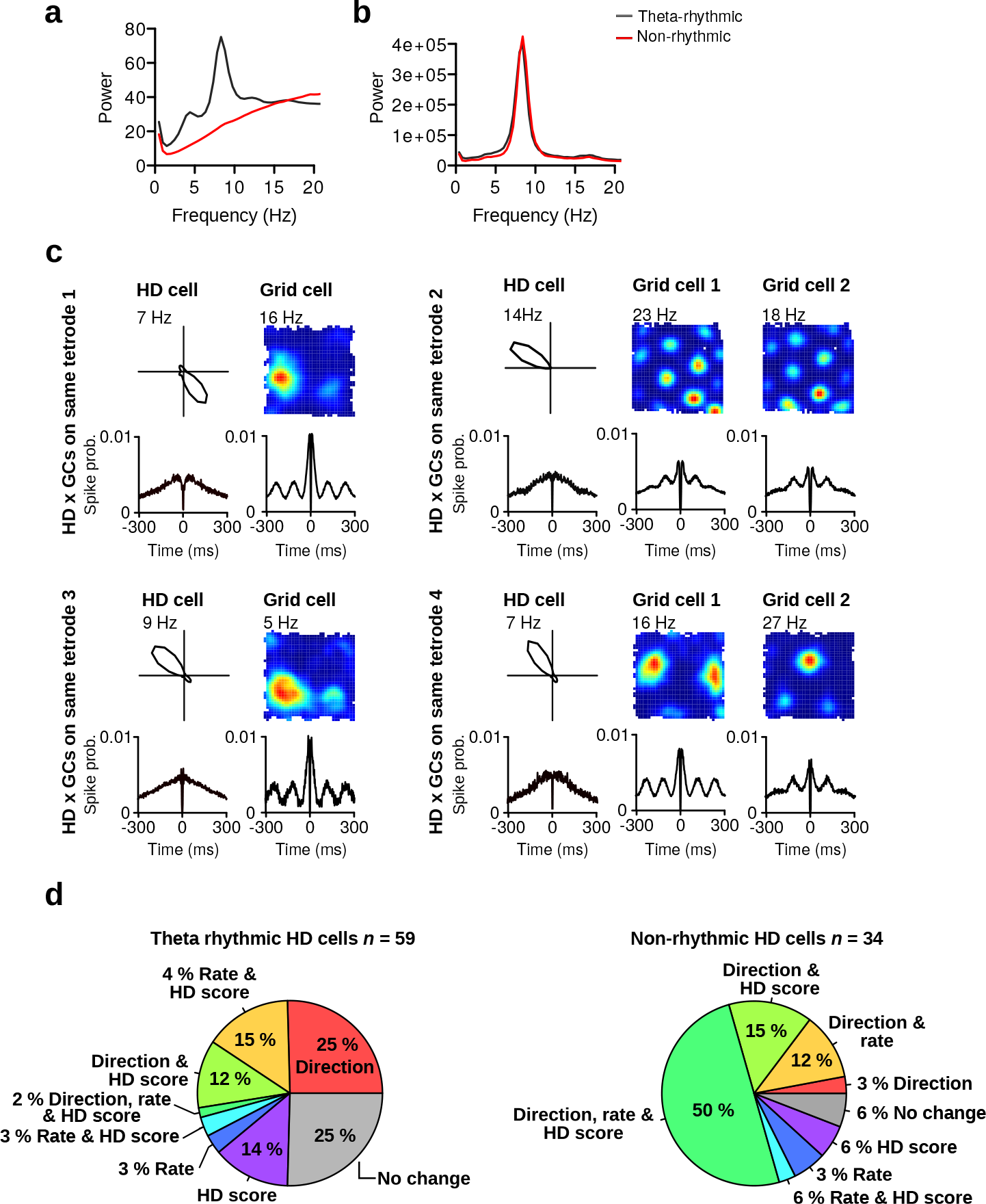
Properties of theta rhythmic and non-rhythmic HD cells. **a**, Average instantaneous firing rate power spectra of theta rhythmic (gray lines) and non-rhythmic HD cells (red lines). **b**, Average local field potential power spectra of theta rhythmic (gray lines) and non-rhythmic HD cells (red lines). **c**, Four examples of non-rhythmic HD cells recorded on the same tetrode as grid cells (GCs). For each HD cell: top row shows the tuning curve of the cell and the spatial firing rate maps of simultaneously recorded grid cells. Bottom row shows spike-time autocorrelations of the same cells. Polar plots and firing rate maps are based on data from vp1 trials. **d**, Percentage of cells with significant changes between vp1 and vp2 trials for theta rhythmic (left) and non-rhythmic HD cells (right).

**Supplementary Figure 4.**
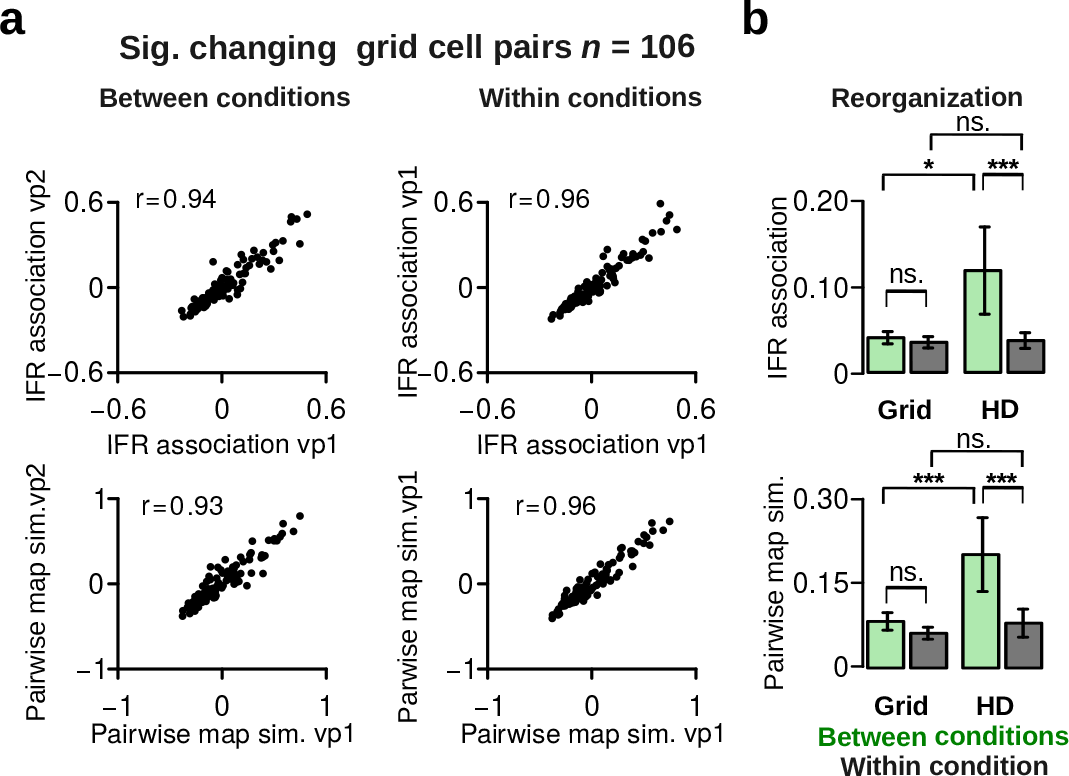
Grid cell pairs with significant visually-driven changes preserve their firing associations. **a**, Comparison of instantaneous firing (IFR) associations (top) and pairwise map similarity of pairs of grid cellls for vp1 and vp2 trials (left) or for two mutually exclusive subsets of vp1 trials (right). *r* values are correlation coefficients. **b**, Reorganization scores of grid cell pairs and HD cell pairs between conditions (vp1 and vp2, gray) or within condition (green). Grid cells significantly changing their map similarity or average firing rate between vp1 and vp2 trials showed no reorganization in their IFR association (top) or pairwise map similarity between conditions (bottom). ns.: not significant, *: *P* < 0.05, ***: *P* < 0.001. Plots show mean ± 95% confidence intervals.

**Supplementary Table 1.** Histological results. Location of the tetrode tips in each hemisphere and number of HD cells (theta rhythmic and non-rhythmic) and grid cells recorded in each hemisphere.

| Mouse | Hemisphere | MEC | PaS/<br>MEC | PaS | PrS/<br>PaS | PrS | PrS/<br>RSA | Cortex | HD<br>cells | Theta-<br>rhythmic | Non-rhythmic | Grid<br>cells |
| --- | --- | --- | --- | --- | --- | --- | --- | --- | --- | --- | --- | --- |
| jp757 | left | 1 | 0 | 0 | 2 | 0 | 0 | 0 | 11 | 4 | 7 | 9 |
| jp757 | right | 1 | 0 | 0 | 0 | 0 | 0 | 0 | 8 | 1 | 7 | 19 |
| jp759 | left | 4 | 0 | 0 | 0 | 0 | 0 | 0 | 0 | 0 | 0 | 13 |
| jp759 | right | 2 | 0 | 0 | 0 | 0 | 0 | 0 | 10 | 3 | 7 | 31 |
| jp865 | left | 0 | 0 | 0 | 0 | 2 | 1 | 0 | 0 | 0 | 0 | 11 |
| jp865 | right | 3 | 0 | 0 | 0 | 0 | 0 | 1 | 7 | 7 | 0 | 2 |
| jp2097 | left | 0 | 0 | 4 | 0 | 0 | 0 | 0 | 2 | 2 | 0 | 1 |
| jp2097 | right | 2 | 0 | 0 | 0 | 0 | 0 | 0 | 17 | 16 | 1 | 8 |
| jp2158 | left | 0 | 0 | 2 | 0 | 0 | 0 | 0 | 0 | 0 | 0 | 0 |
| jp2158 | right | 4 | 0 | 0 | 0 | 0 | 0 | 0 | 2 | 2 | 0 | 23 |
| jp2159 | left | 3 | 0 | 0 | 0 | 0 | 0 | 0 | 4 | 3 | 1 | 30 |
| jp2159 | right | 2 | 0 | 0 | 0 | 0 | 0 | 0 | 8 | 2 | 6 | 33 |
| jp3302 | left | 0 | 0 | 1 | 0 | 0 | 2 | 0 | 0 | 0 | 0 | 0 |
| jp3302 | right | 2 | 0 | 0 | 0 | 0 | 0 | 0 | 8 | 7 | 1 | 0 |
| jp4506 | left | 2 | 2 | 0 | 0 | 0 | 0 | 0 | 6 | 5 | 1 | 20 |
| jp4506 | right | 4 | 0 | 0 | 0 | 0 | 0 | 0 | 9 | 7 | 2 | 11 |
| jp4688 | left | 0 | 0 | 2 | 1 | 0 | 0 | 0 | 0 | 0 | 0 | 7 |
| jp4688 | right | 4 | 0 | 0 | 0 | 0 | 0 | 0 | 1 | 0 | 1 | 1 |
| **Total** |  | **34** | **2** | **9** | **3** | **2** | **3** | **1** | **93** | **59** | **34** | **219** |

